# Network state changes in sensory thalamus represent learned outcomes

**DOI:** 10.1101/2023.08.23.554119

**Authors:** Masashi Hasegawa, Ziyan Huang, Jan Gründemann

## Abstract

Thalamic brain areas play an important role in adaptive behaviors. Nevertheless, the population dynamics of thalamic relays during learning across sensory modalities remain mostly unknown. Using a cross-modal sensory reversal learning paradigm combined with deep brain two-photon calcium imaging of large populations of auditory thalamus (MGB) neurons, we identified that MGB neurons are biased towards reward predictors independent of modality. Additionally, functional classes of MGB neurons aligned with distinct task periods and behavioral outcomes, both dependent and independent of sensory modality. During non-sensory delay periods, MGB ensembles developed coherent neuronal representation as well as distinct co-activity network states reflecting predicted task outcome. These results demonstrate flexible cross-modal ensemble coding in auditory thalamus during adaptive learning and highlight its importance in brain-wide cross-modal computations during complex behavior.

**Summary:** Deep brain imaging reveals flexible network states of sensory thalamus predicting task outcome in mice.

---

Relaying sensory information from subcortical areas to sensory cortices is a fundamental function of thalamus^1,2^, yet we are only starting to understand the computational role of individual sensory thalamic nuclei in flexible encoding during adaptive behaviors^3–7^. Auditory thalamus (medial geniculate body, MGB) flexibly encodes sensory information across associative learning^8–12^, and populations of individual MGB neurons exhibit diverse single cell response adaptations to conditioned tone stimuli in a behavioral state-dependent manner during fear learning^5^. In addition to learning-related auditory response plasticity, MGB neurons process cross-modal sensory inputs in a complex manner^10,13^. For example, visual stimuli enhance MGB tone responses non-linearly^10^, while tactile stimuli affect MGB tone responses bidirectionally^13^, indicating that, similar to cortical and collicular brain areas, MGB processes cross-modal sensory inputs in addition to its auditory relay function^14–17^. These data suggest that MGB is an active computational unit which processes complex information across sensory modalities upon adaptive behaviors. Nevertheless, it remains unknown how large-scale neuronal dynamics in auditory thalamus represent sensory stimuli of different modalities that change their assigned value and expected outcome during flexible learning.

## Results

### Auditory thalamus neurons display various response patterns to cross-modal sensory stimuli in reward associative learning

To address this question, we developed a cross-modal sensory Go/Nogo reversal learning task in mice to test for neuronal population dynamics during sensory learning and for cognitive flexibility upon changing reward contingencies. Mice (N = 8 animals) were trained to associate counterbalanced auditory (12 kHz tone) and visual (rightward drifting grating) stimuli as Go (reward predictor) or Nogo (not rewarded) cues (Fig. 1a-c). Once mice learned the task at expert level (*Initial learning*), the reward contingency was reversed and the previous Go cue became non-predictive of the reward, while the previous Nogo cue turned into a reward-predictor (*Reversal learning)*. Animals learned the initial rule within approximately 10 days (Fig. 1d-f and fig. S1). Upon reversal learning, task performance dropped initially, yet mice learned the new stimulus-reward rule again with a similar learning rate (N = 8 mice, Fig. 1d-g and fig. S1). These data indicate that mice flexibly associate sensory stimuli with reward outcome across initial and reversal learning using similar learning strategies.

**Fig. 1.**
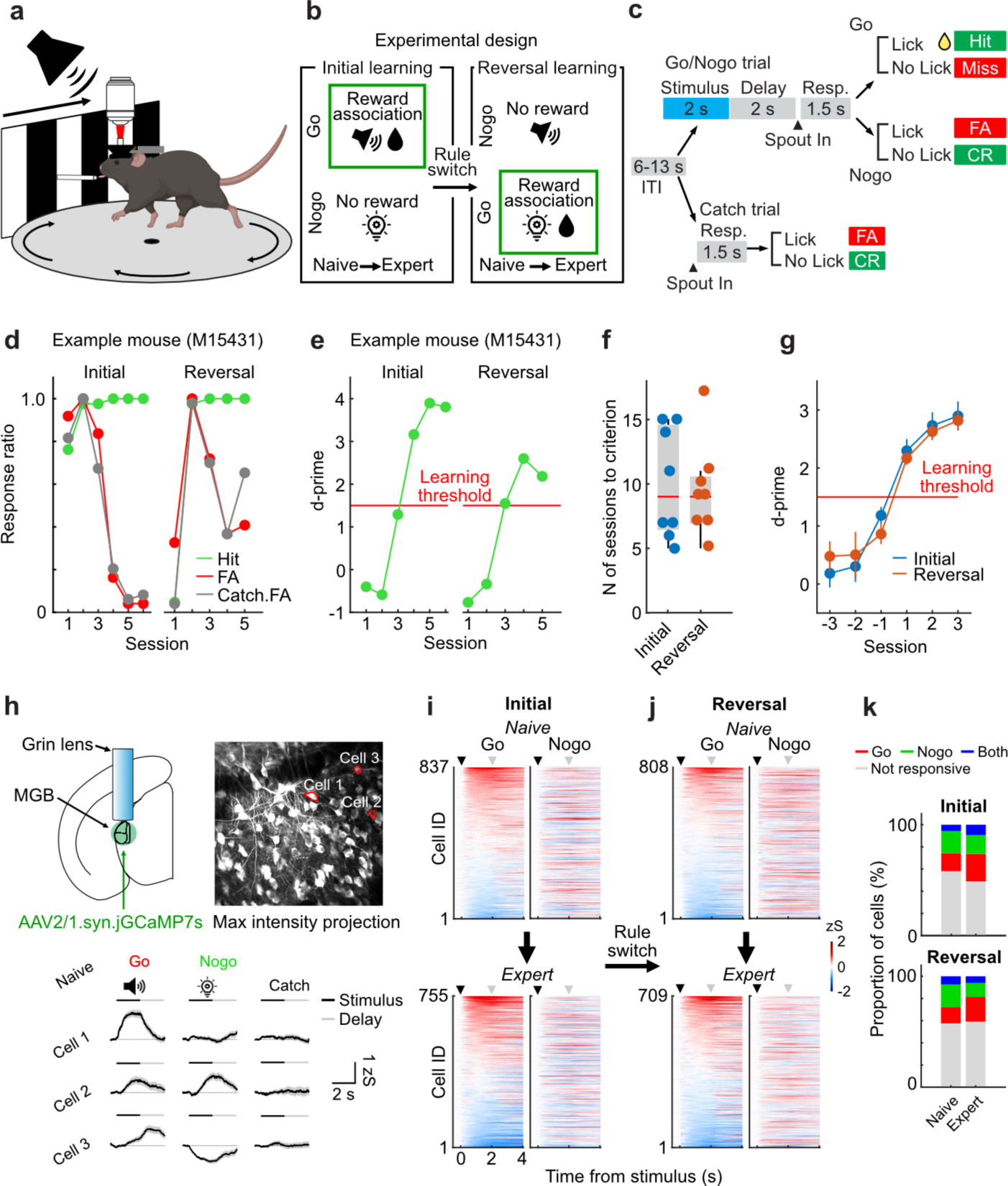
Auditory thalamus neurons exhibit diverse responses and dynamic bias towards reward predictors upon cross-modal reversal learning. **a,** Schematic of the behavioral setup. **b,** Experimental design of the sensory Go/Nogo reversal learning paradigm. **c,** Trial structure. **d,** Task performance of one representative mouse. **e,** Example d-prime transition of the same mouse shown in (d). Learning threshold: d-prime > 1.5. **f,** Number of sessions to reach learning criterion (N = 8 mice). Stimulus-reward association sequence (auditory to visual vs. visual to auditory) is counterbalanced across mice (N = 4 for each group). **g,** Transitions of d-prime around the learning threshold are similar between initial and reversal learning (mean ± SEM, N = 8 mice). **h,** Top left: Schematic of GRIN lens implantation. Top right: Example two-photon image (max intensity projection) of MGB. The contrast is enhanced for improved visualization. Bottom: Ca^2+^ traces from three example neurons in Go, Nogo and Catch trials (mean ± SEM). **i,** Mean individual cell activities in Go and Nogo trials during the initial learning (single session for Naïve and Expert, N = 8 mice). Cells were sorted by the mean response amplitude during the stimulus presentation in Go trials. Cell IDs are matched across Go and Nogo trials in each learning phase. Black and gray triangles represent the stimulus and delay period onset, respectively. **j,** Mean individual cell activities in Go and Nogo trials during the reversal learning (single session for Naive and Expert, N = 8 mice). **k,** Proportion of stimulus-responsive cells from the data shown in **i** and **j**. Top: The transition of the proportion of stimulus-responsive cells from naive to expert phases in the initial learning. The proportions of the stimulus-responsive cells were altered from Naive to Expert phases in both initial and reversal learning (both *p* < 0.01, Chi-square). Parts of panel 1a were created with BioRender.com.

To track the activity patterns of MGB neurons during learning, we performed longitudinal *in vivo* two-photon calcium imaging of large populations of individual MGB relay neurons through a gradient refractive index (GRIN) lens across all stages of the learning paradigm (Fig. 1h, Movie S1). During naive and expert phases of initial and reversal learning, MGB neurons exhibited a large variety of distinct response patterns to the auditory tone as well as the visual stimulus and combinations thereof (Fig. 1h-j, fig. S2), indicating that subsets of MGB neurons are inherently responsive to cross-modal sensory stimuli. Nevertheless, the learning-induced reward association of the Go stimulus during initial or reversal learning altered the proportion of stimulus-responsive MGB neurons towards the Go stimulus regardless of the sensory modality (Fig. 1k, fig. S2). These results demonstrate that auditory thalamus processes cross-modal sensory information during discriminatory reward learning and that MGB responsiveness is dynamically biased towards reward predictors independent of stimulus modality.

### Functional neuronal subgroups predict task-outcome in MGB

Next, we separated MGB neurons into functional subgroups by using a k-means-based cluster analysis approach. MGB neurons exhibited distinct stable, learning-enhanced as well as learning-inhibited responses to the reward-predicting Go stimuli (Fig. 2a,b). These learning-related functional clusters in MGB emerged regardless of the modality of the Go stimulus (Fig. 2a,b, right). In addition to stimulus-driven responses, subsets of MGB neurons developed ramping activity during the non-sensory, pre-reward delay period of Go trials (Ramp-up or Ramp-down clusters), during which the animal has to retain the stimulus type (Go vs. Nogo) and prepare for the action. This ramping activity was specific to the Go stimulus, given that only a small proportion of neurons exhibited ramping activity during Nogo delay periods (fig. S3d,e).

**Fig. 2.**
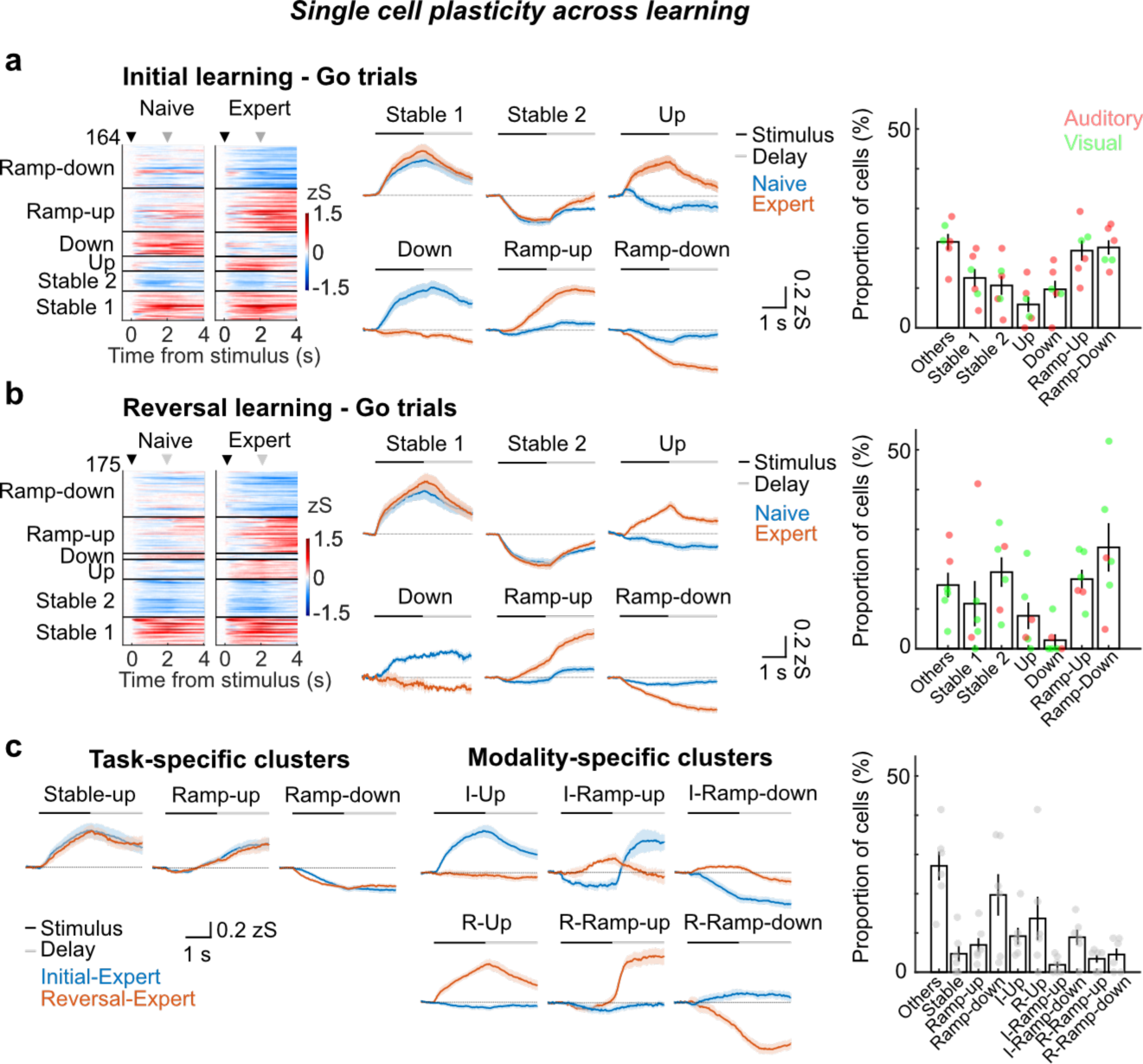
Reward learning induces heterogeneous single cell plasticity in a task- and modality-specific fashion. **a,** Functional subgroups of MGB neurons in initial learning. Left: Heatmaps of single cell activities before and after learning in initial learning. Cells were clustered into functional subgroups depending on their activity patterns (n = 164 cells from 6 mice). Middle: Average calcium traces (mean ± SEM) of the functional subgroups shown on the left side. Right: Proportion of cells in each cluster. Each dot represents the data from the individual mouse. Red and green dots represent the type of stimulus-reward association. **b,** Learning-related functional subgroups in reversal learning (N = 6 mice, n = 175 cells). Figure structures are the same as (a). **c,** Functional subgroups exhibiting task and modality-specific plasticity in the two expert phases in initial and reversal learning. Left: Average calcium traces (mean ± SEM) of the task-specific functional subgroups. Middle: Average calcium traces (mean ± SEM) of the modality-specific functional subgroups. Right: Proportion of cells in each cluster.

We then compared the activity patterns of the same neurons across expert states in initial and reversal learning when the animals are successfully performing the task, and found task-specific MGB neurons that are active during distinct trial epochs irrespective of the sensory modality (Fig. 2c, left). In contrast, a second modality-specific group of MGB neurons was modulated by the trial epoch in a sensory modality-specific manner (Fig. 2c, middle). These results demonstrate that reward learning drives heterogeneous neuronal plasticity in MGB that flexibly reflects task features and reward outcomes in a modality-driven as well as modality-independent manner.

### Functional plasticity in MGB is not exclusively driven by behavioral variables

During the cross-modal Go/Nogo paradigm, mice adapted their behavior flexibly and reversibly. Behavioral variables such as body movement, licking^9,18,19^ or arousal level^20^ affect neural activity in a brain-wide manner. Depending on the trial epoch, approximately 5.7-38.6 % of MGB neurons exhibited correlations of Ca^2+^ activity with behavioral variables (fig. S4a,b, threshold: r = 0.2). However, the behavioral modulation of Ca^2+^ activity was not systematically changed across learning (fig. S4c,d). In addition, we found that the strength of the correlation of Ca^2+^ activity and locomotion or pupil size in neurons that were tracked across the behavioral paradigm was unchanged or even decreased after learning and upon behavioral adaptation (fig. S4e,f). In addition to locomotion and arousal, the number of licks during the pre-reward delay period increased after learning (fig. S4g,h). Nevertheless, anticipatory licking did not systematically modulate MGB activity during expert phases (fig. S4i,j) and did not correlate with the proportion of ramping cells during the delay period (fig. S4 k). Taken together, these results indicate that subsets of individual MGB neurons correlate with behavioral variables such as locomotion, pupil size and licking, yet changes in behavior are not the main driver of functional plasticity and changes in learning-related activity in MGB.

### Learning induces coherent neuronal population representations during reward-preceding periods

In addition to the classification of individual stable and plastic MGB neurons, we next asked if and how the neural population representation of MGB changes upon learning. Throughout an individual session as well as across learning, the trial-by-trial population vector correlation (PVC, Pearson’s r)^21–23^ of the stimulus period (Go and Nogo trials) remained stable (Fig. 3a,b, figs. S5-6). In contrast, the PVC specifically increased during the reward-preceding delay period of Go trials once mice learned the stimulus-reward contingency in initial learning and flexibly adapted to the previous non-rewarded delay period after reversal learning (Fig. 3c,d, figs. S5-6). Thus, upon associative learning, the neuronal representation of the MGB population becomes more similar during the reward-preceding delay period in a sensory modality-independent fashion (figs. S5-6). The PVC increase was specific to the outcome and not the action given that it occurred only during the delay period of Hit, but not False Alarm trials, where mice incorrectly licked to a Nogo stimulus. (Fig. 3e,f, and figs. S7-8). Furthermore, the delay period PVC increase was present in trials with and without anticipatory licking (Fig. 3g) indicating that enhanced PVCs during the delay period are not driven by the preparatory behavior of the animal or lick impulsivity. Removing delay cells (i.e., Ramp-up and Ramp-down cells, see Fig. 2a,b; see fig. S9 for the removal of unspecific subclusters) did not affect the increase in PVC (Fig. 3h), indicating that the change in population representation during the delay period is driven by the total population of MGB neurons.

**Fig. 3.**
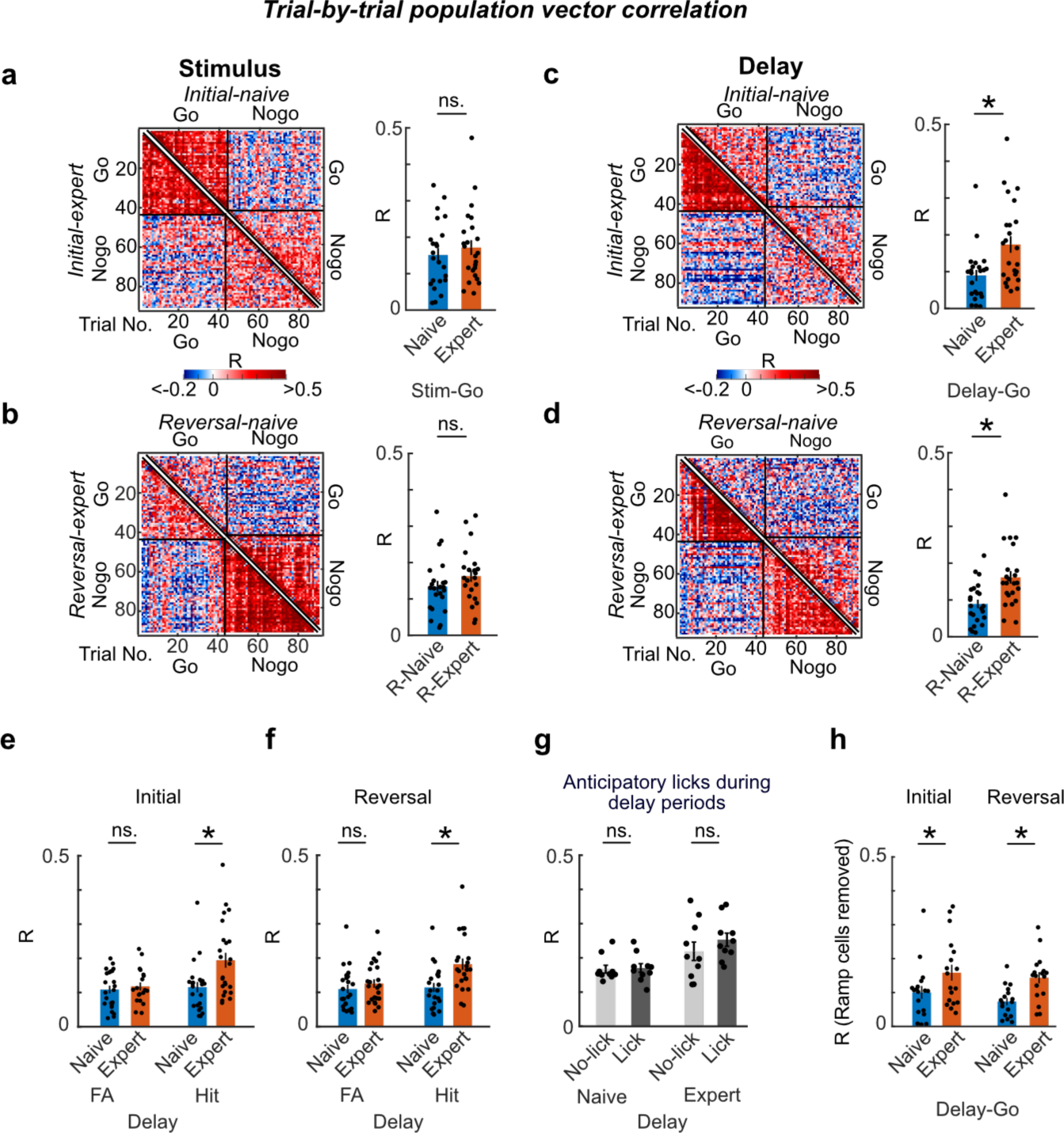
Learning-induced remodeling of neural population level representations of task features in MGB. **a-d,** Representative single session trial-by-trial population vector correlation matrices of stimulus (a, b) and delay (c, d) periods from one example mouse. Bar charts show the means of Go trial PV correlation between naive and expert phases in initial (a, c) and reversal learning (b, d) from all mice (N = 8 mice, 2-3 sessions / mouse, see figs. S5-8). **e,f,** Mean trial-by-trial population vectors correlation of the delay period of Hit and False Alarm (FA) trials in different learning stages (N = 8 mice, 1-3 sessions / mouse, see figs. S7, 8). **g**, Mean trial-by-trial correlation in the delay period in Hit trials with or without anticipatory licking in the initial learning (N = 8 mice, 1-3 sessions / mouse, see figs. S5, 6). Due to the small number of the Hit trials with licking in reversal learning, we excluded the reversal learning data for this analysis. **h,** Mean trial-by-trial correlation in the delay period after removing all ramping cells found in Fig. 2a,b in all mice (N = 6 mice, 2-3 sessions / mouse, see fig. S9). Each dot is the mean R value in individual sessions in each training stage for all mice. Statistical tests: Linear mix model (a-f, h, *p < 0.01, details in Table S1). Wilcoxon rank sum test (g).

### Associative reward learning changes the co-activity network structure in MGB

Next, we analyzed how the co-activity network structure changes across learning in MGB by computing the weighted undirected graphs of MGB population activity^24,25^. Here, nodes in the co-activity network represent individual MGB neurons and edges the positive pairwise Pearson’s correlation coefficients of neural activity between all neuron pairs (Fig. 4a). During the delay periods of the Go trials, the hubness (average of the activity correlations between all cell pairs) increased from the naive to the expert phases, while it was unaffected in the Nogo trials (Fig. 4b). Furthermore, the mean shortest path length between any two neurons became shorter in the expert phase in the Go trials, which is consistent with a global increase of hubness (Fig. 4b). Local co-activity structures represented by the cluster coefficient of triad did not increase from the naive to expert phases (fig. S10a), indicating that the observed changes are a global event in the total MGB population. Upon reversal learning, the co-activity structure during the delay period was actively remodeled, both globally (hubness and path length) and locally (cluster coefficient) (Fig. 4b, fig. S10a), indicating that MGB co-activity network structure is dynamic during changing stimulus-reward contingencies. Furthermore, removing the cluster of delay ramping neurons (or a similar number of random neurons) from the analysis did not affect the changes in the co-activity network structure (fig. S10b,c). These results indicated that the general MGB population mediates the remodeling of the co-activity network. Anti-correlation networks (negative Pearson’s r) remained unchanged in all learning stages (fig. S11). In summary, global co-activity network structures in MGB changed dynamically during the reward-preceding delay period and flexibly re-adjusted after learning in a modality-unspecific fashion. Changes in co-activity structures during the stimulus period were less consistent and limited to the reversal learning period (fig. S12). Altogether, these results demonstrate that the functional co-activity structure in MGB can be rapidly re-organized through associative reward learning and learning rule switches, which could help to support the cognitive processing of task-relevant information^24,25^.

**Fig. 4.**
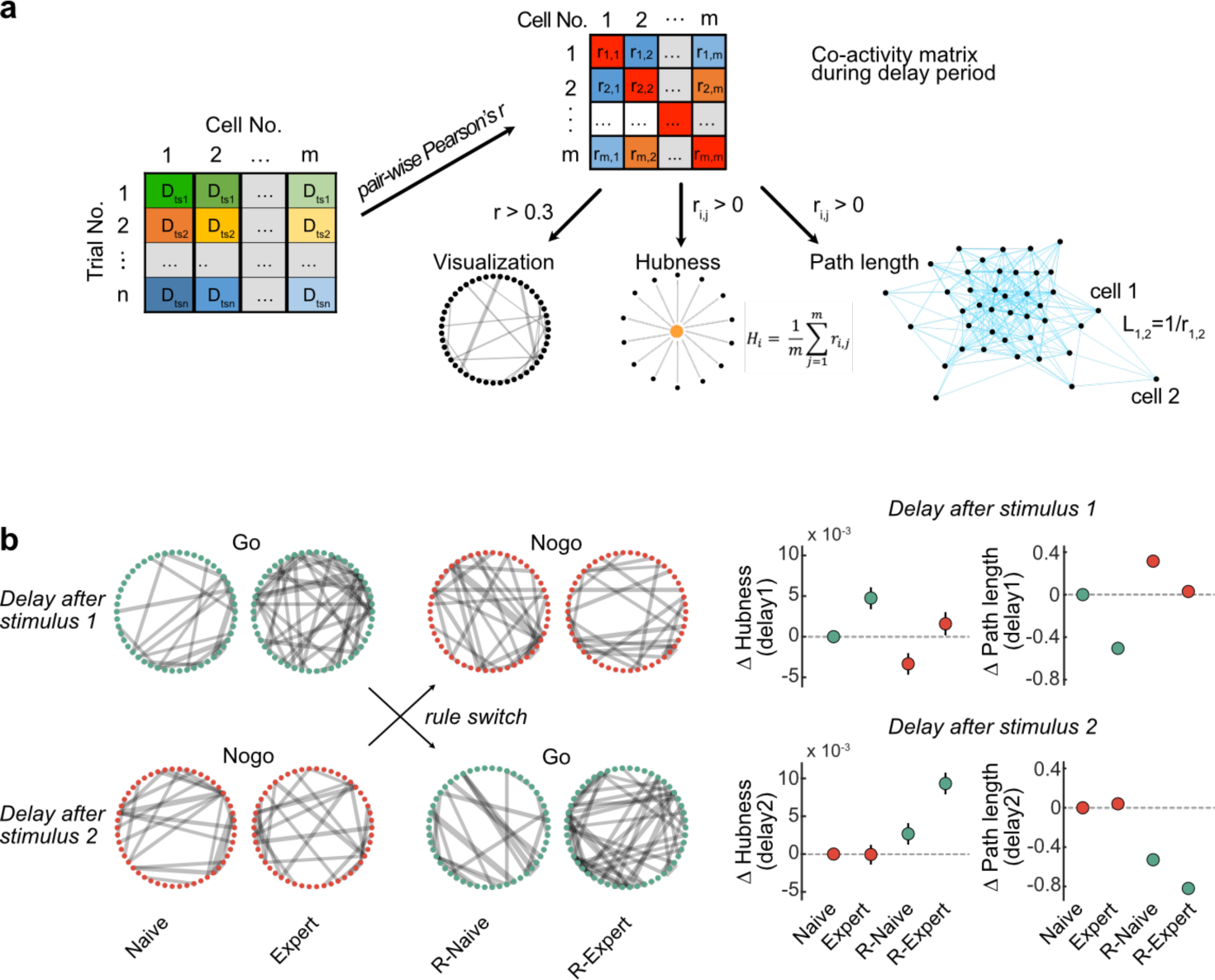
Learning re-organizes MGB co-activity structure. **a,** Co-activity matrix construction and network analysis. The co-activity matrix represents the cell-by-cell pair-wise Pearson’s correlation (r) from all concatenated time series vectors of the delay periods (D_ts_) from all trials (see left matrix for construction of time series vector). The co-activity strength is visualized by line thickness in a circular plot for each cell pair (r > 0.3, dots indicate individual neurons). Global MGB co-activity strength is quantified by hubness and shortest path length. D_tsn_: 2 s time series during delay period in trial n. r_i,j_: Pearson’s correlation of calcium activity between cell i and j. H_i_ : hubness of cell i to all its neighbors. L_1,2_: path length between cell 1 and 2. **b,** Left: Representative circular plots show an example MGB co-activity network structure at different learning stages of Go and Nogo trials for one representative mouse before and after learning the rule switch. Right: Changes of hubness and path length across all neuron pairs in each learning stage which are baselined to the values in the naive Go (Stimulus 1) or Nogo (Stimulus 2) condition (N = 210 neurons from 6 mice, 3000 bootstraps). Error bars: 95 % confidence interval of mean.

## Discussion

Recent work has shed light on the role of sensory thalamus in adaptive behaviors^3,4^, while the neural dynamics of sensory thalamus in flexible learning of complex tasks across multiple sensory modalities remained elusive. Here, we combined a cross-modal (auditory and visual) reversal learning paradigm with longitudinal deep-brain two-photon calcium imaging in medial geniculate body (MGB) and demonstrated that sensory thalamus dynamically encodes sensory as well as task-related information, adapting its neural responses to varying task rules.

We find that auditory thalamus exhibits adaptive processing of auditory and visual information (Fig. 1h-j and figs. S13, 14) similar to sensory cortices^15,16^. Responses of MGB neurons to auditory and visual stimuli are plastic and modulated upon cross-modal stimulus-reward learning (Fig. 1k), indicating that sensory encoding in MGB is dynamic upon cognitively demanding tasks irrespective of the sensory modality (fig. S2). A subset of MGB neurons developed ramping activity during the reward-preceding delay period (Fig. 2a-c), during which the animal has to hold the predictive information of the stimulus about the trial outcome, which could reflect reward anticipation^11^ or short-term memory to guide the next action^26^, similar to cortical and thalamic areas of the mouse motor system^27^. Reciprocal loops between motor thalamus and cortex have been shown to be necessary to maintain ramping activities in either region during delay periods^27^, suggesting that, similar to the motor system, auditory thalamus could cooperatively maintain delay ramping activity together with auditory and other cortices.

On the population level, learning can either increase^28,29^ or decrease^30,31^ trial-by-trial correlation, whilst other cognitive factors such as attention^32,33^ exhibit a bi-directional influence. In our reversal learning task, the neuronal representation of sensory stimuli in auditory thalamus was stable across learning, regardless of whether they were predictive of a reward (Go cue) or not (Nogo cue). These findings are similar to previous observations for conditioned stimulus presentations in aversive auditory fear learning^5^. In contrast to the stimulus period, the MGB population response (trial-by-trial correlation) became more coherent across trials during the non-sensory pre-reward delay period, once mice learned the task-reward rule, irrespective of the sensory modality (Fig. 3a-d, figs. S5, 6). This trial-by-trial population vector correlation increase during the delay period was specific to Hit trials, but not False Alarm trials (Fig. 3e, f, figs. S6, 7), indicating that the re-organization of the population activity during the delay period is dependent on the predicted task outcome and not the general reward preparation and consumption movement per se (e.g., licking, Fig. 3g). Recently, trial-by-trial correlations were suggested to account for the optimization of information communication between upstream sensory information encoding and downstream read-out guiding behaviors in different population representation subspaces^34–36^. Together, our findings link changes in population dynamics of auditory thalamus to a wider brain network that is implicated in functions of task representation^37^, short term memory^27,38^, outcome-dependent actions^39^ and reward anticipation^11^.

Co-activity network structure during the reward-preceding delay period was remodeled across associative learning regardless of the sensory modality, while it remained by-and-large stable during the stimulus period (Fig. 4b, figs. S10, 12). The distinct changes of co-activity network states between the stimulus and delay periods may result in differential information transfer to downstream areas to prime information integration and guide behavioral outputs in Go or Nogo trials^7,40,41^. While the global co-activity network changed upon learning (i.e., path length and hubness), the local co-activity network (i.e., cluster coefficient) did not systematically change during the delay period, and ramping cells did not contribute to the enhancement of the co-activity network (fig. S10). This suggests that the activity of the total MGB population supports the functional network structure and not individual functional subgroups of neurons. Upon reversal learning, the co-activity structures were dynamically remodeled not only during the delay but also during the stimulus periods (fig. S12), which indicates that reversal learning might depend on a distinct learning rule, resulting in the generalization of the Go stimulus from a sensory representation towards a predictive state representation that could be flexibly assigned to future learning rule updates. It remains to be tested if and how these changes are mediated and whether they are generated locally or if they are co-dependent on external activity, e.g., primary or associative cortical brain areas^42^.

What are the neural mechanisms that guide the functional plasticity in MGB from naive to expert phases across reward associative learning? Given the sparse local recurrent connectivity within mouse sensory thalamus^1,43,44^ and negligible proportions of local inhibitory neurons^5,45^, local microcircuit plasticity is unlikely to drive changes in response patterns. Plausible scenarios could include changes of synaptic plasticity, neuromodulation or adaptations of long range bottom-up excitatory^46,47^ and/or inhibitory^3,4^ inputs. In addition, corticothalamic feedback^2,48^ could stabilize changes in MGB population activity upon learning, either directly or di-synaptically via the thalamic reticular nucleus (TRN). Future studies will need to test how distinct circuit elements and their combination can affect functional population level plasticity in auditory thalamus.

While reward learning biased the sensory responses of MGB neurons towards reward-associated Go stimuli during the task regardless of sensory modality (Fig. 1k, fig. S2), the sensory responses in off-task non-rewarded mapping sessions before and after learning (i.e., passive measurements of tuning to multimodal sensory stimuli) were not biased to the on-task Go stimuli (figs. S13, 14). Specific multisensory enhancement to the Go stimulus, which was previously observed during appetitive learning in MGB^10^, did not take place during off-task, non-rewarded sessions (figs. S15, 16). This could be due to a fast devaluation of reward-conditioned stimuli^11^. Finally, while uni and multisensory responses could be altered on the single cell level (figs. S13, 14), sensory stimuli could be reliably decoded from the MGB population activity across learning (fig. S17). The complementary mechanisms of single cell plasticity and population-level stability of sensory coding could be crucial to allow for dynamic neuronal representations upon learning, while ensuring stable representations of the environment^5,49^.

Altogether, our study reveals that auditory thalamus displays flexible adaptations of single cell responses and co-activity network states that align not only with sensory but also task-period and outcome-relevant information, which change bi-directionally upon reversal learning, highlighting the role of sensory thalamus in complex neural computations for adaptive behaviors.

## Supporting information

Source Data Main

## Acknowledgments

We thank the HHMI Janelia GENIE Project for making jGCaMP7 available; Raymond Strittmatter, Patrick Schlenker, Robert Häring and Simon Saner for mechanical workshop and electrical engineering support; all members of Gründemann laboratory for their general support and feedback on the manuscript.

## Funding

Swiss National Science Foundation (SNSF professorship PP00P3_170672, J.G.)

European Research Council Starting Grant 803870 (J.G.)

The Forschungsfonds Nachwuchsforschende of the University of Basel (M.H.)

Forschungspreis der Schweizerischen Hirnliga (J.G.)

The Department of Biomedicine at the University of Basel (J.G.)

Deutsche Forschungsgemeinschaft (DFG), SFB 1089 (Teilprojekt, J.G.)

Deutsches Zentrum für Neurodegenerative Erkrankungen (DZNE), Bonn, Germany (J.G.)

## Author contributions

Conceptualization: MH, JG

Methodology: MH, ZH, JG

Investigation: MH, ZH, JG

Visualization: ZH, MH, JG

Funding acquisition: JG, MH

Project administration: JG, MH

Supervision: JG

Writing – original draft, review & editing: MH, ZH, JG

## Data and materials availability

Data to generate figures and supplementary figures in this paper can be accessed in Data S1 and S2, respectively. Statistical results are shown in Table S1. Custom-written code used to generate figures and supplementary figures are available at https://gin.g-node.org/GrundemannLab/ThalamicNetworkStates_Code. Data is available at https://gin.g-node.org/GrundemannLab/ThalamicNetworkStates_Data.

## Methods

### Animals

All experiments were performed in accordance with the institutional guidelines of University of Basel and were approved by the Cantonal Veterinary Office of Basel-Stadt, Switzerland. Six to 10-week-old (at the start of the experiment) male C57BL/6J mice were used throughout this study. Animals were housed on a 12-hour light / dark cycle at an ambient temperature (22 ° C) and humidity (55 %) and had free access to food and water until the initiation of the behavioral experiment. Throughout the behavioral experiment, animals were placed under food restriction and their body weights were maintained at 85-90% of their free-feeding weights. Well-being was monitored daily through the entire experimental period. No statistical methods were applied to pre-determine the sample size for each experiment. The investigator was not blinded for surgery, behavioral and imaging experiments or data analysis.

### Surgical procedures

Buprenorphine (0.1 mg/kg) was subcutaneously injected approximately 30 min before surgery for analgesia. Then, mice were anesthetized with isoflurane (1.5 - 2.0% maintenance) through the oxygen-enriched air (95 %, 1-3 l/min, Oxymat III, Weinmann). Anesthesia level was monitored via breathing rates and foot and tail reflexes before and during surgery. Mice were placed in a stereotaxic apparatus (Model 1900, Kopf Instruments), and their body temperature was maintained through a heating pad (Rodent warmer, 53800M, Stoelting). Their eyes were covered by an eye protective cream (Bepanthen Augen und Nasensalbe, Bayer). The mixture of Lidocaine (10 mg/kg) and Ropivacaine (3 mg/kg) was injected under the skin over the skull for local anesthesia. Stereotaxic viral injections were performed as previously described^5^. Briefly, a small craniotomy was performed above the medial geniculate body (MGB, AP: −3.28, ML: −1.9, DV: −3.0 mm) by using a stereotaxic drill (Model 1911, Kopf) with a burr drill bit (105-0135-225, Kyocera). A pulled glass pipette (2-000-001, Drummond Scientific) filled with AAV vector was slowly lowered into the brain with the help of a micropositioner (Model 2650, Kopf). AAV2/1.syn.jGCaMP7s ^50^ (Addgene, 104487-AAV1, ca 500nl, diluted by sterile PBS, 1-2x) was injected into MGB with a pressure ejection system (Picospritzer, Parker). One to two weeks after viral injection, a gradient refractive index (GRIN) lens (0.6 mm diameter, 7.3 mm length or 1.0 mm diameter, 4 mm length, Inscopix) was implanted during the second surgery (anesthesia and analgesia, see above). 0.6 mm lenses were implanted as previously described ^5^. Briefly, a 0.8 mm diameter craniotomy was performed above MGB (drill: 105-0709.400, Kyocera) and a small track was cut with a 0.7 mm sterile needle. Next, the GRIN lens was slowly advanced into the brain using the Micropositioner (Model 2650). For the implantation of 1.0 mm diameter lenses, a 1.2 - 1.3 mm craniotomy was performed above MGB using a hand drill (503599, World Precision Instrument) with a burr drill bit (200 µm diameter, C1.104.002, Bösch Dental). Tissue above MGB was slowly aspirated through a sterile blunt needle (27G, Endo irrigation cannula) connected to a suction system. Sterilized phosphate buffer saline (PBS) was used to irrigate the brain until the bleeding stopped around the aspirated site. Next, the 1.0 mm lens was slowly advanced into the brain with the micropositioner. Both, 0.6 and 1.0 mm lenses were fixed to the skull with light curable glue (Loctite 4305, Henkel). A custom-made head bar was attached to the skull next to the GRIN lens, and the skull was sealed with Scotchbond (3M), Vetbond (3M) and dental acrylic (Paladur, Kulzer, Orth Jet, Lang Dental and/or C&B Super-Bond, Sun Medical). Meloxicam (5 mg/kg) was injected subcutaneously after the surgery for post-operative analgesia.

### Behavioral apparatus

The behavioral apparatus was housed in a light shield chamber under a custom-built two-photon microscope (Independent NeuroScience Services (INSS), UK). Mice were head-fixed by a custom-designed holding system and placed on a running-wheel connected to a rotary encoder (E6A2-CS3E, Omron) to measure locomotion. Auditory and visual stimuli were presented with a speaker (ES1, Tucker Davis Technologies, placed at upper-right, 10 cm from the mouse head) and a 7-inch screen (Adafruit 1667, placed 10 cm from the right side of the mouse face at a parallel angle), respectively. The screen system was modified to minimize the light exposure to photomultiplier tubes (PMTs) during a visual stimulus presentation in the two-photon imaging ^51,52^. During the experiment, the gray background was continuously presented from the screen. A lick spout was mounted on the custom-built retractable stage controlled by Trinamic motion control language (TMCL). A reward (soy milk) was delivered by a custom-designed peristaltic pump system by using a micropump (mp6, Bartels Mikrotechnik). A licking of a reward spout was detected by a lick detector modified from the detector described in a previous study ^53^. The experiments were controlled by a custom-written program in MATLAB (MathWorks) with NI USB-6008 (National Instruments) and RZ 6 (Tucker Davis Technologies), and the timing of TTL input/output of behavioral events were recorded by RZ 6 at 50 kHz sampling rate.

### *In vivo* two-photon calcium imaging

*In vivo* two-photon calcium imaging was performed using a custom-built two-photon microscope (INSS, UK). The microscope was equipped with a resonant scanning system and a pulsed Ti:sapphire laser (λ = 940 nm, Chameleon Vision S, Coherent). A motorized three-axis system (Zaber motor) connected with a microscope head (Z-direction) and breadboard under the behavioral apparatus (X-Y directions) enabled locating an objective lens above the GRIN lens. The microscope system was controlled by ScanImage software (Vidrio Technologies). Green and red fluorescent photons were collected with an objective lens (x16, 0.80 NA, Nikon). Photons were separated by a dichroic mirror (T565lpxr, long pass, Chroma) and barrier filters (green: ET510/80m, red: ER630/75m), and measured by PMTs (PMT2101, Thorlab). The imaging frame was 512 x 512 pixels, and the frame rate was approximately 30 Hz. Fields of views (FOVs) of two-photon images were approximately 330 µm x 330 µm (at 3.0x zoom, 1.0 mm diameter GRIN lens, N = 6 mice) or approximately 400 µm x 400 µm (at 2.5x zoom, 0.6 mm diameter GRIN lens, N = 2 mice). Note that GRIN lens FOVs do not correspond to the actual size of the imaged brain areas due to the spatially non-uniform optical distortion inherent to GRIN lenses^54^.

### Two-photon Image Processing

Two-photon images were processed using Suite2P^55^. The images were motion-corrected, and regions of interest (ROIs) were automatically generated. Next, experimenters curated ROIs and sorted them as neurons or not. A small portion of ROIs were drawn using a manual ROI drawing function of Suite2p. A neuropil signal was also calculated for each ROI by Suite2P. A correction coefficient (0.7) was multiplied with the neuropil signal, and each ROI signal was subtracted from this value and handled as Ca^2+^ signal of individual neurons. To extract the same ROI signals through the behavioral and sensory mapping sessions, we employed ROIMatchpub (https://github.com/ransona/ROIMatchPub). For behavioral sessions, we selected 2-3 sessions per learning phase (initial naive, initial expert, reversal naive and reversal expert) for each mouse. The naive phase consisted of the first 2-3 behavioral sessions, and the expert phase consisted of the 2-3 sessions in which d-prime (d’, discriminability index)^56^ reached over 1.5. Across these selected sessions, we tracked the same ROIs wherever possible. The ROI tracking procedure was done separately for behavioral and sensory mapping sessions. The same ROI signals, or matched cell data, were used for the following analyses: k-means clustering of the Go/Nogo task (Fig. 2a-c, fig. S3), co-activity network structure analysis (Fig. 4 and figs. S9-12), correlation analysis for behavioral variables and Ca^2+^ signals (fig. S4a-f) as well as the sensory mapping analysis (figs. S13-17) unless stated otherwise. For the trial-by-trial population vector correlation analysis, co-activity network structure analysis as well as decoder training (see below), Ca^2+^ signals of individual neurons were detrended and lowpass filtered to 5 Hz with a Butterworth filter.

### Sensory mapping

Before the start of the sensory mapping sessions, mice were head-fixed under the custom-built two-photon microscope and habituated to the behavioral apparatus and environment for minimum 3 days. Each session started with a 1 min habituation period. Auditory (4, 8, 12, 16 or 20 kHz pure tones at 75 dB, 2 s), visual (Upward, downward, rightward or leftward sine wave drifting gratings, 100% contrast, 2 Hz, 0.05 cycle per degree, 2 s) or multisensory stimuli (combination of auditory and visual stimuli, e.g., 4 kHz pure tone with upward drifting grating, 2 s) were presented. The sensory stimuli were presented in a pseudo-random order. Each auditory, visual and the multisensory stimulus was presented 8 times (e.g., 4 kHz pure tone, rightward grating, and the combination of them were presented for 8 times), and a total of 240 trials (8 trials x 30 stimulus types of uni and multisensory stimuli) were performed in one session per day. Inter-trial interval (ITI) was 6-9 s. For one mouse, each stimulus was presented 5 times and a total of 150 trials / session were performed. Sensory mapping was performed for 2-3 consecutive sessions before and after the sensory Go/Nogo reversal learning paradigm.

### Behavioral training in a sensory Go/Nogo reversal learning task

Following sensory mapping sessions, mice were pre-trained to lick a spout to receive a liquid reward (soy milk) for 1-2 days under head-fixation. Next, the animals were trained to perform a sensory Go/Nogo task, which consisted of Go trials (30%), Nogo trials (35%) and catch trials (35%) (total number of trials: 140 per session). At the beginning of each trial, a 6-13 s ITI was initiated. Then, either an auditory stimulus (Go cue, 12 kHz pure tone, 75 dB, 2 s), a visual stimulus (Nogo cue, rightward drifting grating, 2s) or no stimulus (2 s blank period, catch trial) was presented. The sensory stimulus or blank period was followed by a delay period (2 s) without any sensory stimulus. At the end of the delay period, a response window (1.5 s) was initiated and a retractable lick spout moved forward to the mouse. Go trials required mice to lick the spout (Hit) to obtain a reward (8-10 µl soya milk), otherwise the trial was considered as an error (Miss). In Nogo trials, mice were required to withhold the lick response (Correct Rejection, CR). If the mice licked the spout in the Nogo trial, the trial was considered as an error (False Alarm, FA). In catch trials mice were required to withhold the lick response (catch-CR). If mice licked the spout, the trial was considered as an error (catch-FA). If mice accidentally touched the spout too early (within 200 ms after the response window onset) (e.g., due to grooming), the contact was not considered as a response. Catch trials were introduced to ensure that mice identify the Go cue as a reward predictor. Task performances for Nogo and catch trials developed similarly (Fig. 1d,e, and fig. S1). Thus, the task performances in those trials were combined to calculate the task performance index, d-prime (d’) as follows: d’ = Z (Hit ratio) - Z (False alarm ratio). Hit and False alarm ratios were calculated as follows: Hit ratio = the number of Hit trials/(the number of Hit trials + the number of Miss trials); False alarm ratio = the number of false alarm trials/(the number of false alarm trials + the number of correction rejection trials). If the ratio reached 1.0, the ratio was adjusted to calculate d’ ^56^. Z is the inverse cumulative distribution function, and Z (Hit ratio) and Z (False alarm ratio) were calculated by using the qnorm function (https://de.mathworks.com/matlabcentral/fileexchange/48978-qnorm-matprobabilities-dblmean-matsigmas-booluseapproximation). Once d’ reached values above 1.5 for three consecutive days, mice were considered experts and a reversal learning paradigm was initiated. Upon reversal learning, the stimulus-reward contingency was switched. In the group in which the auditory stimulus was presented as a Go cue, the visual stimulus was now presented as a Go cue and rewarded and vice versa for animals in which the visual stimulus was initially presented as a Go cue. Once d’ reached values above 1.5 for three consecutive days, mice were considered reversal experts. The order of the stimulus-reward contingency was counterbalanced between mice, i.e., four mice were trained to the auditory stimulus as a Go cue first and four mice were trained to the visual stimulus as a Go cue first. Specific cases of data exclusion: Mice were trained for a maximum of 28 sessions. Due to slower learning, one out of eight mice reached only two reversal expert sessions before the training had to be terminated. Furthermore, in two out of eight mice the training had to be briefly suspended due to technical issues, which did not have a major impact on task performance after re-initiation of the paradigm. In one mouse, an imaging artifact gradually appeared from the lateral edges of the two-photon image in one session. Data from the later stage of this session was excluded from analysis. In one mouse, the data recording of one session was aborted in the middle of the session due to the malfunction of the acquisition system. Since the number of trials reached more than half of the session, the data obtained in this session was included for the data analyses.

### Video recording

Mouse behavior was monitored during the experiment with a CMOS camera (a2A2590-60umPRO, Basler) equipped with a CCTV lens (Moritex ML-M1616UR) and a band-pass filter (DB850, Midwest) (N = 7 mice) or a Raspberry Pi Camera Module 2 with a shortpass filter (FES850, Thorlabs) controlled by Raspberry Pi 3 model B+ (N = 1 mouse). Either camera system was located on the left side of the mouse together with a custom-made infrared LED system (830 nm). In the CMOS camera system, each frame acquisition was synchronized with the acquisition of a two-photon image (ca. 30 Hz) through ScanImage. In the raspberry Pi camera system, each frame was acquired at ca. 30 Hz in free-run mode and image acquisition was monitored by internally-generated TTLs. During offline analysis, the tongue and pupil were detected and tracked by animal pose estimation using DeepLabCut (DLC)^57^. The experimenter manually labeled the tongue and the eight points on the edges of the pupil (top, top-right, right, bottom-right, bottom, bottom-left, left, top-left) for each mouse to train the model. Videography based licking-behavior was detected if the tongue was tracked by DLC for at least two consecutive frames. The pupil area was calculated by using a circle fit function^58^ (fitcircle, https://mathworks.com/matlabcentral/fileexchange/15060-fitcircle-m?s_cid=ME_prod_FX) from available data points at each frame. Pupil area data was smoothed by using a hampel filter (MATLAB built-in function, number of neighbors, 10; number of standard deviations, 1.0).

### Mean individual cell activity during Go and Nogo trials

Heatmaps were used to visualize mean activities of individual MGB neurons pooled from all mice across learning (N = 8 mice, Fig. 1i, j). For the naive phase, the data of the first training session in both initial and reversal learning were used to show the unconditioned responses to sensory stimuli. For the expert phase, the data from the session with the highest d’ in both initial and reversal learning were used to show well-conditioned responses to the sensory stimuli. The calcium data was baselined to the mean during 0.5 s before stimulus presentation in each trial. The data of individual calcium traces were averaged across Go and Nogo trials separately (maximum 42 trials for Go trials, and maximum 49 trials for Nogo trials in a single session). Cell IDs were sorted according to the amplitude of the mean sensory response during the stimulus presentation (2 s) in the Go trials.

### Proportion of stimulus-responsive cells in the sensory Go/Nogo task

Stimulus-responsiveness was determined through a two-step procedure. First, we performed a signed-rank test to examine if the sensory response of each cell was significantly different from zero. In each cell, the sensory response during the stimulus presentation (2 s) was averaged in Go and Nogo trials. Then, the means of the sensory responses pooled across Go or Nogo trials were used for a signed-rank test of each cell. Cells with statistical significance in the signed-rank test were selected for response thresholding (threshold: median z-score > ± 0.2). In addition, for auditory stimulus trials, the sensory response during the stimulus onset (0.3 s) was averaged and analyzed in the same manner as described above to catch fast-adapting MGB neurons. If a neuron was classified as sensory responsive during the whole 2 s stimulus presentation period or as onset responsive, the neuron was included as auditory responsive. Neurons where then classified as Go, Nogo or ‘both’ responsive cells depending on the trial type. Neurons that did not pass the detection threshold were classified as non-responsive cells.

### K-means cluster analysis

K-means cluster analysis was performed to sort individual neurons into functional subgroups. The calcium traces of matched individual neurons (N = 6 mice, n = 210 cells) were averaged across Go or Nogo trials for 2-3 sessions in each training phase (naive, expert, reversal naive and reversal expert). The mean Go or Nogo responses of each neuron were concatenated between naive and expert phases in both initial and reversal learning (time-series concatenation). The time-series for each cell was composed of the concatenated stimulus and delay periods of native and expert training phases, while initial and reversal learning were treated as independent observations. This times series underwent principal component analysis followed by k-means clustering (cosine distance) to sort the individual neurons into functional clusters. Thirty clusters were generated, and clusters showing similar activity patterns were merged manually ^5^. After generating the merged clusters, cell IDs in each merged cluster were separated to the initial and reversal learning data. To track the activity patterns of the MGB neurons between the two expert phases in initial and reversal learning, we performed k-means cluster analysis by using the data of the two expert phases in the initial and reversal learning (Fig. 2c, fig. S3a-c). After the preprocessing described above, the data of the two expert phases were concatenated between initial and reversal learning, then k-means clustering was performed.

### Correlation analysis between behavioral variables and neuronal activity

Correlations between behavioral variables (locomotion, pupil size) and single cell Ca^2+^ traces were calculated as the Pearson’s correlation coefficient in Hit trials across learning (fig. S4a-f). The first session in the naive phase and the session with the highest d-prime in the expert phase were selected in both initial and reversal learning. Pupil, locomotion and calcium data of day-matched cells was down-sampled to 5 Hz. Cells that exhibited correlations between the behavioral variable and the calcium data of |r| > 0.2 and p < 0.05 were considered as significantly correlated. R-value distributions were compared between the naive and expert phase in both initial and reversal learning by a Kolmogorov-Smirnov test (ks-test2, MATLAB). To measure the change of correlation across learning, the difference of |r| values between naive and expert phases in matched cells was measured for stimulus, delay and ITI periods (signed-rank test). The pupil data was analyzed in the same manner except that the pupil data was smoothed by a Hampel filter (see above) and z-scored (fig. S4b,d,f).

### Anticipatory licking analysis

Anticipatory licking was quantified during the delay period. To visualize how the number of anticipatory licks change across learning, the proportion of trials with anticipatory licking was plotted as a function of the number of anticipatory licks per trial. The number of trials with or without anticipatory licking was pooled from all mice in each training phase, and this pooled data was used for chi-square testing (fig. S4g). The probability of anticipatory licks per time bin was calculated across mice (fig. S4h). For visualization purposes, lick probability was smoothed by a moving average (5 frames, movmean, MATLAB). To examine how anticipatory licking influenced MGB activity during the delay period, Ca^2+^ traces of Hit trials with and without anticipatory licking were separated and baselined to the mean Ca^2+^ fluorescence 150 ms before the delay period onset (fig. S4i, same baseline duration for fig. S4j).

### Trial-by-trial population vector correlation

Before the construction of the population vector (PV), calcium data for each cell was baselined to the mean of the 2 s pre-stimulus period on a trial-by-trial basis. Next, the mean calcium responses during the stimulus and delay periods were calculated and the population vector constructed. Trial-by-trial population vector correlations (PVC, Pearson’s) were calculated across the stimulus and delay periods of all trials and plotted as a N-by-N correlation matrix per session (N = number of trials). Mean correlation values from all trial pairs in each session were used for summary statistics. A linear mixed model was used to compare the difference of R values (fixed effect: naive vs expert, random effect: mouse ID). For a fair comparison between trials with and without anticipatory licking, only sessions with more than 6 trials with and without anticipatory licking each were included in the analysis (Fig. 3g). Reversal learning was excluded from the analysis of anticipatory licking due to low trial numbers. To examine the contribution of ramping cells to the PVC during the Go delay period ramp-up and ramp-down cells were removed from the PV, and PVC values of the Go delay period were re-calculated. The same number of cells as ramp-up and ramp-down cells were randomly removed to generate the shuffle dataset (nShuffle = 30, fig. S9).

### Weighted graph-based analysis of MGB co-activity network structure

Undirected weighted graph that represents the relationship of calcium activity between MGB neurons were computed based on matched cell data (n = 210 neurons). In the graph, nodes and edges stand for cell identities and the pairwise Pearson’s correlation coefficients between two neuron pairs, respectively. The calcium activity data was baselined to the mean of the 2 s pre-stimulus period of each trial. The pair-wise correlation coefficient of the time-series data (2 s) during the stimulus and delay periods was calculated between all cell pairs in Go and Nogo trials. Each graph with the number of nodes M, which are identical to the number of matched cells is represented by its adjacency M-by-M symmetric matrix C where each element r_ij_ is the Pearson’s correlation coefficients (−1≤ r ≤1) between two nodes (neurons) i and j. Positive and negative edges were analyzed separately (fig. S11). The strength of the global co-activity network structure among MGB neurons was quantified by hubness, i.e., the mean of the pair-wise activity correlation of each node to all the others:

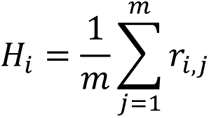

We estimated the global communication efficiency between any two nodes by measuring the geodesic (shortest) path length in the graph. First, a weighted graph with the length between two nodes, i and j as L_ij_=1/r_ij_ was created. Next, the shortest path length between any two nodes was computed using Dijkstra’s algorithm among 3903 cell pairs. The local co-activity network structures were quantified by the cluster coefficient of the triad. The cluster coefficient of the triad was calculated among 50376 triads ^59^. w_ij_ is the edge weight between any two nodes. *ŵ*_ij_ is the normalized edge weight by the maximum weight in the adjacency matrix between cell i and j:

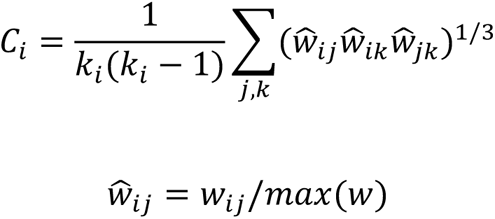

The same number of cells as ramp-up and ramp-down cells were randomly removed to generate the shuffle dataset (nShuffle = 30, fig. S10). Network structure was recalculated using the shuffle dataset. Functional connectivity parameters were calculated using the BrainConnectivity toolbox (https://github.com/jblocher/matlab-network-utilities/tree/master/BrainConnectivity). Plotting and confidence interval calculation were adapted from DABEST toolbox (https://github.com/ACCLAB/DABEST-Matlab).

### General sensory responsiveness

To quantify the general sensory responsiveness of MGB neurons before and after the Go/Nogo learning in sensory mapping session (fig. S13), the calcium data during the stimulus presentation (2 s) was split in half (0 - 1 s and 1 - 2 s). The mean of the calcium data during these two periods was handled as individual data points. Next, it was tested whether individual MGB neurons were responsive to auditory, visual and/or multi-sensory stimulus through the same two-step procedures as described above. In multi-sensory trials, the data from all visual grating directions were averaged, and the sensory response to auditory frequencies were tested. In fig. S13c, cells were defined as excited/inhibited if they were responsive to any auditory frequency or grating direction. In fig. S13d, the response amplitude of the cells that were responsive to at least one of the sensory stimuli in pre-learning were compared to the response amplitude of the same sensory stimuli in post-learning sessions. In fig. S13e, change of peak tuning frequency or grating direction was calculated if a cell was responsive to any frequency/direction in both pre- and post-learning session. In fig. S13g and h, the time series of the averaged population response and the response amplitude of the cells with the significant response to the reward-associated stimulus in Go/Nogo training (12 kHz, rightward drifting grating, 12 kHz with rightward drifting grating) were compared between the pre- and post-learning session. In fig. S13i, the change of peak tuning frequency or grating direction was calculated if a cell was responsive to the conditioned stimulus (12 kHz, rightward-drifting grating, 12 kHz with rightward drifting grating) both in pre- and post-learning session.

### K-means clustering of tuning curves

To sort MGB neurons into groups with similar frequency tuning patterns, k-mean clustering (‘correlation’ distance) was performed with 20 features (5 frequencies, multi-/uni-sensory and pre/post training) (fig. S15). Sensory responses to the auditory (4, 8, 12, 16 and 20 kHz) and multi-sensory stimuli (4, 8, 12, 16 and 20 kHz with drifting gratings) were averaged across trials and sessions (1-3 sessions). Visual stimulus feature (i.e., grating directions) was collapsed in multi-sensory trials. The calcium data in the response period was baselined to 0.5 s before stimulus presentation. The mean of the calcium activity during the stimulus presentation (2 s) was used to generate frequency tuning curves. F: mean calcium response during stimulus presentation period. m: cell number (pooled across 6 mice). f: auditory frequency. Clustering matrix:

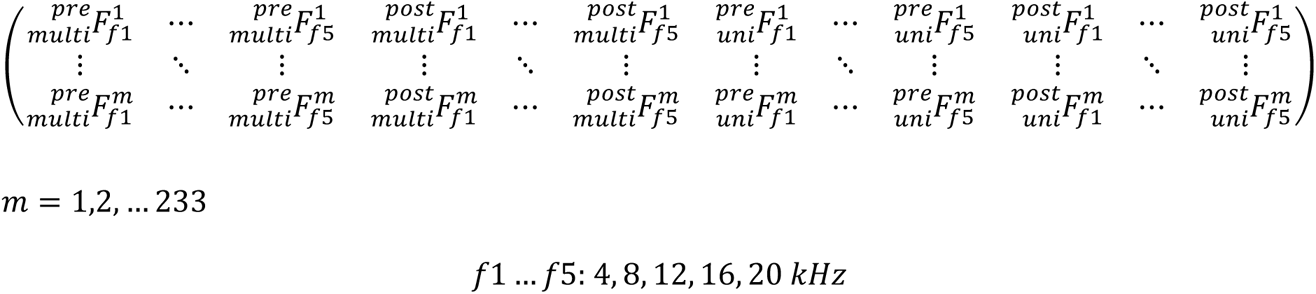

To examine if frequency tuning curves were stable or plastic, similarity of the frequency tuning curves between pre- and post-learning sessions for each cell was quantified by Pearson’s correlation (R) in auditory and multi-sensory trials separately. The distribution of Pearson’s R pooled across all cells was compared between auditory and multi-sensory trials by ks-test (fig. S15c) to examine in which trial the frequency tuning curve would be more plastic or stable.

### Multi-sensory index

The multi-sensory index was calculated by the division of the average response in multi-sensory trials (AV) to the sum of average response in uni-sensory trials (A: auditory trials, V: visual trials). Cells with a non-significant response in both AV and A trials were excluded from multi-sensory index calculation. Cells with different calcium response sign in multi-sensory response and sum of uni-sensory response were excluded (*A*_*f*_*V* ⋅ (*A*_*f*_ + *V*) < 0). Absolute value of sensory response was taken to demonstrate the principle of multi-sensory integration (linear or non-linear). Visual grating directions were averaged in multi-sensory response as well as uni-sensory response. First order exponential fitting was conducted to reveal the trend of data distribution for each auditory frequency (fig. S15d). Multi-sensory indexes of all individual cells and all auditory frequencies were compared regardless of response sign and amplitude (fig. S16). Two-dimensional two-sample ks-test was performed to test if the distribution of the multi-sensory index was comparable between pre- and post-learning (See, Table S1).

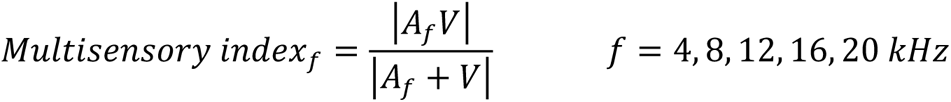

### Single cell correlation analysis across multisensory mapping sessions

Single cell averaged peri-stimulus time histograms (PSTH) of uni- and multisensory trials for stimulus features (auditory frequency, grating direction) in pre- and post-learning were sorted by stimulus amplitude (0 – 2 s) and plotted as heat maps (fig. S14). Next, the Pearson’s correlation between the response time series of individual neurons before and after learning was calculated and average across all cells (fig. S14)^60^.

### Decoder analyses

Linear support vector machine (SVM) decoders were trained on the stimulus period (normalized by 1 s pre-stimulus baseline) for all tracked neurons in each multi-sensory mapping session for each animal. Time series data were down-sampled to 3 Hz to avoid overfitting. The decoders trained on individual sessions were tested in a pair-wise manner for all sessions (fig. S17a). For modality decoding (fig. S17b), decoders were trained on 70 % of the data and tested on a 30 % hold-out test-set. Same numbers of trials from each modality were selected for training and testing in 50 iterations to avoid a biased accuracy measurement due to unbalanced trial number in each modality (50 % multi-sensory, 16.7 % auditory, 13.3 % visual). For auditory frequency decoding (fig. S17c), trial sub-sampling was not necessary since each auditory frequency was presented for a same number of trials. Uni-sensory and multi-sensory trials were combined in one dataset. Visual stimulus feature was collapsed in multi-sensory trials leaving only frequency labels (4, 8, 12, 16 and 20 kHz). Train/test ratio remained 70/30 (140 training trials, 60 test trials). Simple accuracy (modality: x / 27 trials, frequency: x / 60 trials, x = number of correct decoded trials) in all iterations were averaged yielding pair-wise session decoding accuracy. Shuffle data (n = 100) were generated by circular permutation of the down-sampled time series features for each train/test iteration (n = 50) from which mean accuracy values of shuffle iterations were taken to depict chance decoding accuracy and further averaged to yield pair-wise session decoding accuracy (fig. S17b,c).

### Histology

After training, mice were transcardially perfused with phosphate buffer saline (PBS) followed by ca. 40 ml 4 % paraformaldehyde (PFA) in PBS. Immediately after perfusion, brains were removed and post-fixed in 4 % PFA overnight at 4 °C. Then, brains were stored in PBS at 4 °C until dissection. 150 µm coronal slices were prepared using a vibratome (Campden Instruments) and immunostained for calretinin as an anatomical marker as described previously ^5^. Briefly, after PBS washes, brain slices were immersed in blocking solution (10% normal horse serum, S-2000-20, Vector Laboratories) with 0.5 % Triton (T8787, Sigma-Aldrich) in PBS for 2 hours at room temperature. Next, slices were incubated in primary antibody (goat anti-calretinin, 1:1000, CG1, Swant) in carrier solution (1% normal horse serum with 0.5% Triton PBS) overnight at 4 °C. Slices were washed again in 0.5% Triton PBS and incubated for 2 h at room temperature or overnight at 4 °C in secondary antibody (donkey anti-goat 647, 1:1000, A-21447, ThermoFisher) in carrier solution. After final washes by PBS, slices were mounted on slides and cover slipped using 22 x 50 mm, 0.16 - 0.19 mm thick cover glass (FisherScientific). Images were acquired with a LSM700 confocal microscope (Zeiss), Axio Scan Slide Scanner (Zeiss) or Olympus BX63. Acquired images were post-processed with ImageJ (https://imagej.nih.gov/ij/) to locate the implantation site of the GRIN lenses.

### Statistical methods

Statistical analysis was performed in MATLAB (MathWorks). Alpha level was set at 0.05 and Bonferroni correction was applied to statistical tests (see Table S1). Chi-square test, rank sum test, signed-rank test, linear mixed model (LMM), two-sample Kolmogorov-Smirnov test (ks-test2) and two-dimensional two-sample Kolmogorov-Smirnov test (https://github.com/brian-lau/multdist) were performed for datasets indicated in Table S1. Data are presented as mean ± SEM unless otherwise stated. Statistical results and p values are presented in Table S1.

## Movie S1

An example movie of Hit trials in naïve and expert phases. For visualization, the images are cropped and resized. The movie of the calcium imaging was also temporally smoothed.

## Data S1

Source data for Figures in main text (Figs. 1-4).

## Data S2

Source data for Supplementary Figures.

---

**fig. S1.**
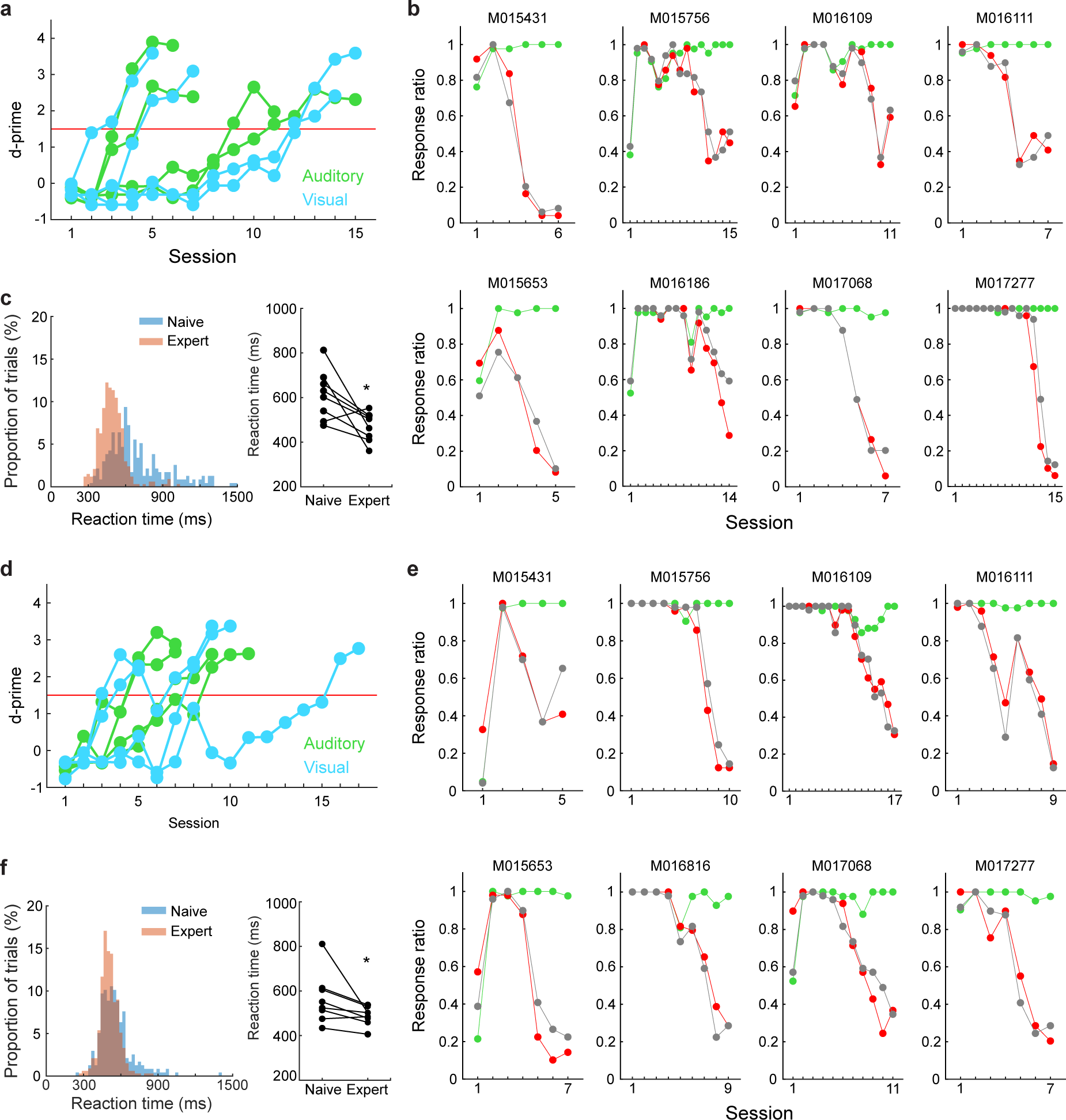
Task performance of individual mice during the sensory Go/Nogo task. (**a-c**) The task performance of individual mice and Hit reaction time (RT) in the initial learning. (**a**) Transition of d-prime from all mice. (**b**) Transition of the task performance of individual mice. Top row: Auditory-reward group. Bottom row: Visual-reward group. (**c**) Left: Hit RT pooled across all mice in the naive and expert phases. Right: Transition of median Hit reaction time. Each dot represents the individual mouse data. RT in the expert phase was shorter than that in the naive phase (signed-rank test). The first behavioral session in the naive phase and the session with the highest d-prime in the expert phase were used for the RT analysis. (**d-e**) The task performance of individual mice and Hit RT in the reversal learning. The figure configurations are the same as **a**-**c**. Reaction time in the expert phase was shorter than that in the naive phase (signed-rank test). * in the figures represent the statistical significance. *, *p* <0.05.

**fig. S2.**
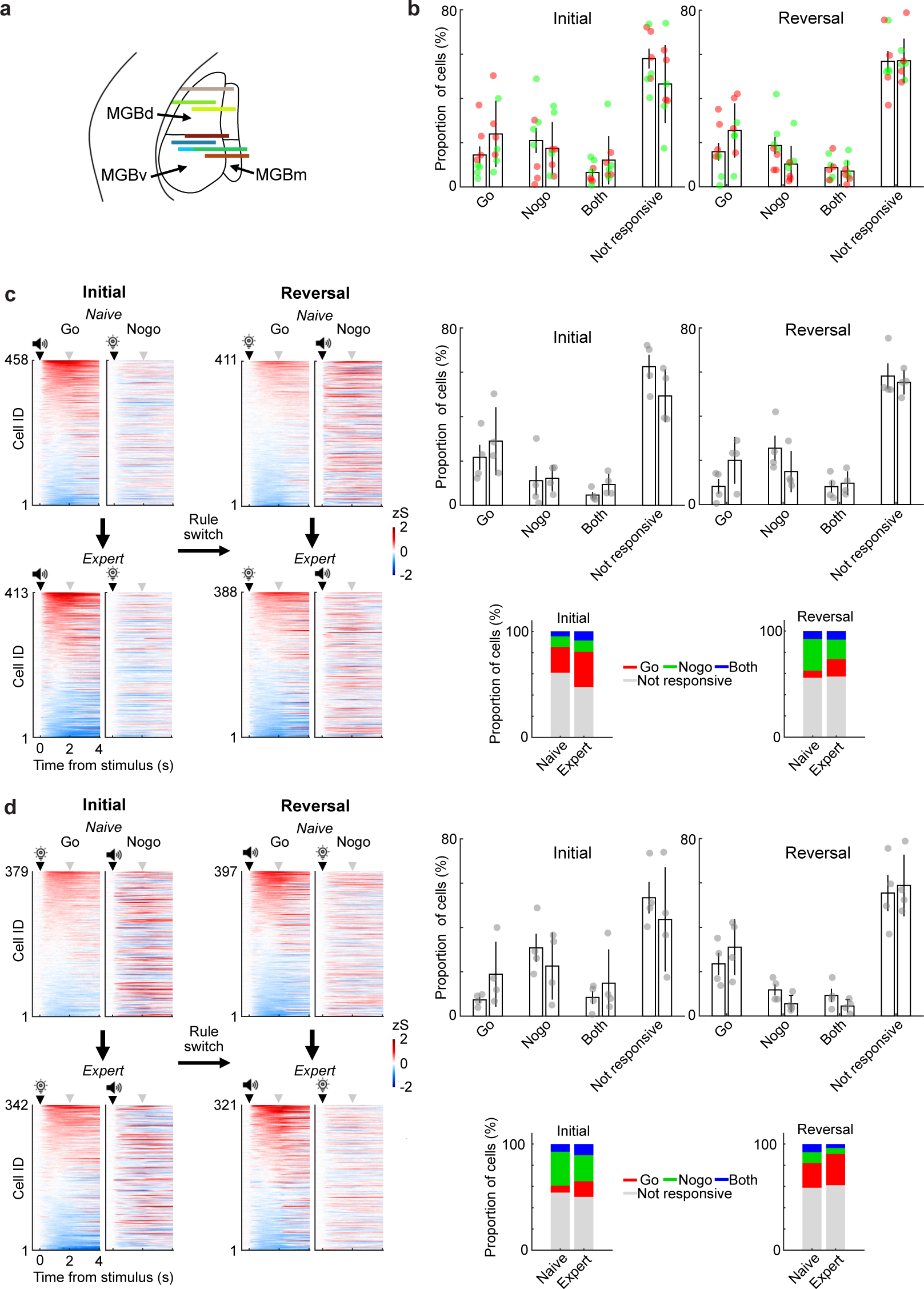
Activity of individual cells and the proportion of stimulus-responsive cells in auditory and visual reward groups. (**a**) GRIN lens front and putative field of view (FOV) from all mice (N = 8 mice). Each line represents the lens front and the putative FOVs of the two-photon image. For the 1.0 mm lens, the FOV is approximately 330 um x 330 um. For the 0.6 mm lens, the FOV is approximately 400 um x 400 um. Due to the distortion of the image through the GRIN lens, these FOV values do not necessarily reflect the actual size of the brain image. (**b**) The proportion of stimulus-responsive cells from all mice (N = 8 mice). (**c**) Mean individual cell responses and the proportion of stimulus-responsive cells from Auditory-reward (initial learning) → Visual reward (reversal learning) group. Left: Heatmaps of the mean individual cell activities in Go and Nogo trials of a single representative session during the initial learning (n = 458 cells in the naive phase, n = 411 cells in the expert phase cells from 4 mice) and reversal learning (n = 413 cells in the naïve phase, n = 388 cells in the expert phase from 4 mice). Cells were sorted by the averaged response amplitude during the stimulus presentation in Go trials. Cell IDs are matched across Go and Nogo trials in each learning phase. Black and gray triangles represent the stimulus and delay period onset, respectively. Right top: The proportion of stimulus-responsive cells in initial and reversal learning. Right bottom: The stacked-bar charts of the stimulus-responsive cells from naive to expert phase in the initial and reversal learning. The proportions of stimulus responsive cells was altered from the naive to the expert phase across learning in both initial and reversal learning (*p* < 0.01, chi-square test, in both initial and reversal learning). (**d**) Mean individual cell responses and the proportion of stimulus-responsive cells from Visual-reward (initial learning) → Auditory reward (reversal learning) group. The figure configurations are the same as (**c**). The proportions of stimulus responsive cells were altered from the naive to the expert phase across learning in both initial and reversal learning (*p* <0.01 for Initial and *p* < 0.05 reversal learning, after Bonferroni correction, chi-square test).

**fig. S3.**
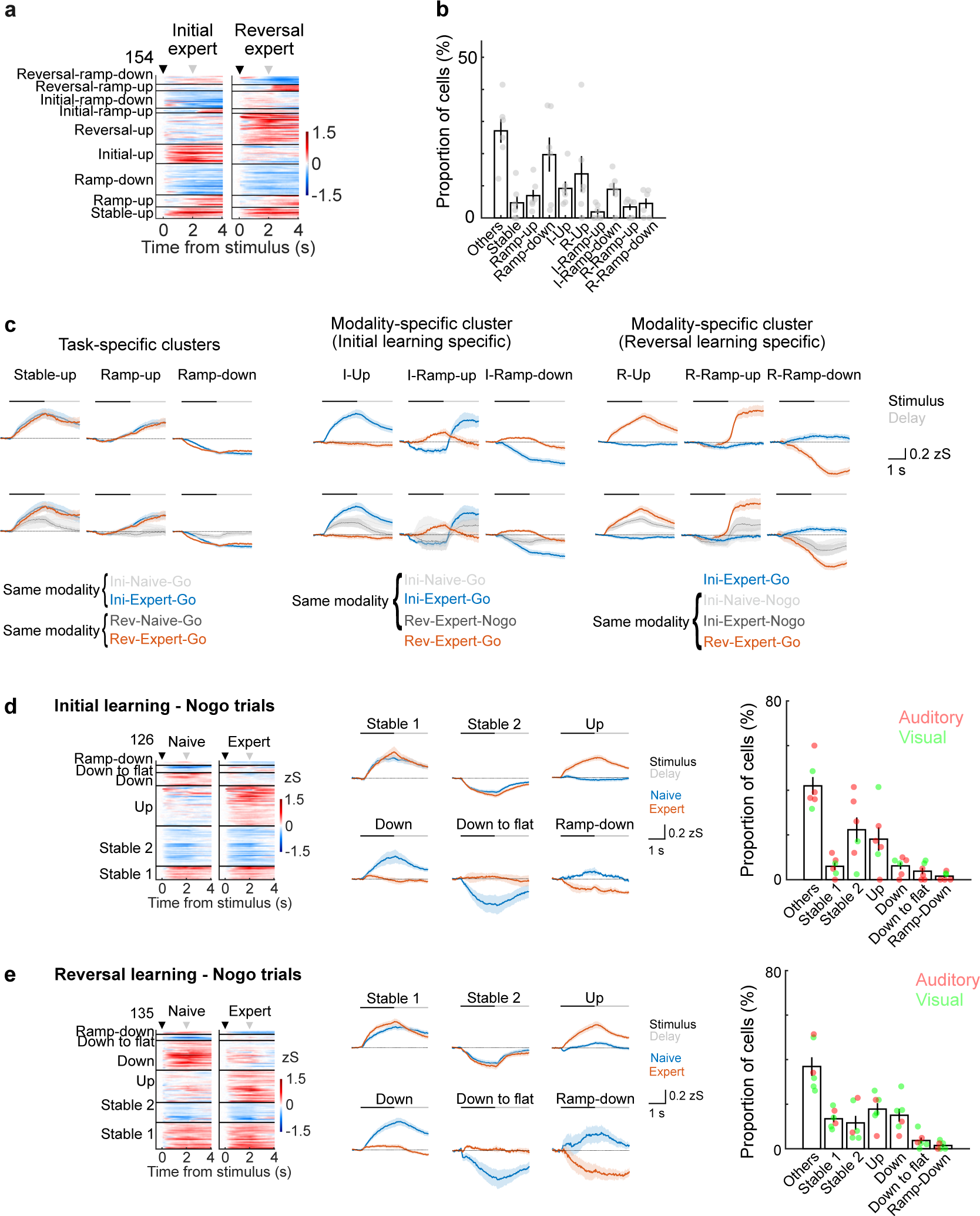
Task-specific and modality-specific clusters in Go trials and functional clusters in Nogo trials. (**a-c**) Functional subgroups with the task and modality-specific learning plasticity in the two expert phases in both initial and reversal learning. (**a**) Heatmaps of single cell activities in the two expert phases (N = 6 mice, n = 154 cells). (**b**) Proportion of cells in each functional cluster. Each dot represents the data from the individual mouse. (**c**) Average calcium traces (mean +/− SEM) of the functional subgroups shown in (**a**). Top row: Average calcium traces in the two expert phases. Left: Task-specific clusters. Middle: Modality-specific clusters showing the plasticity in the initial learning. Right: Modality-specific clusters showing the plasticity in the reversal learning. Bottom row: Average calcium traces of the two expert phases with additional traces of other conditions with the same sensory modality. The additional traces were overlaid to show how reward learning modulated the neuronal activity within the same sensory modality. The figure configuration from Left to Right is the same as the top row. (**d**) Functional subgroups of MGB neurons in Nogo trials in initial learning. Left: Heatmaps of single cell activities in the naive and expert phases in initial learning (N = 6 mice, n = 126 cells). Middle: Average calcium traces (mean +/− SEM) of the functional subgroups. Right: Proportion of cells in each cluster. Each dot represents the data from the individual mouse. Red and green dots represent the types of stimulus-reward condition. (**e**) Functional subgroups of MGB neurons in Nogo trials in reversal learning (N = 6 mice, n = 135 cells). The figure structures are the same as (**d**).

**fig. S4.**
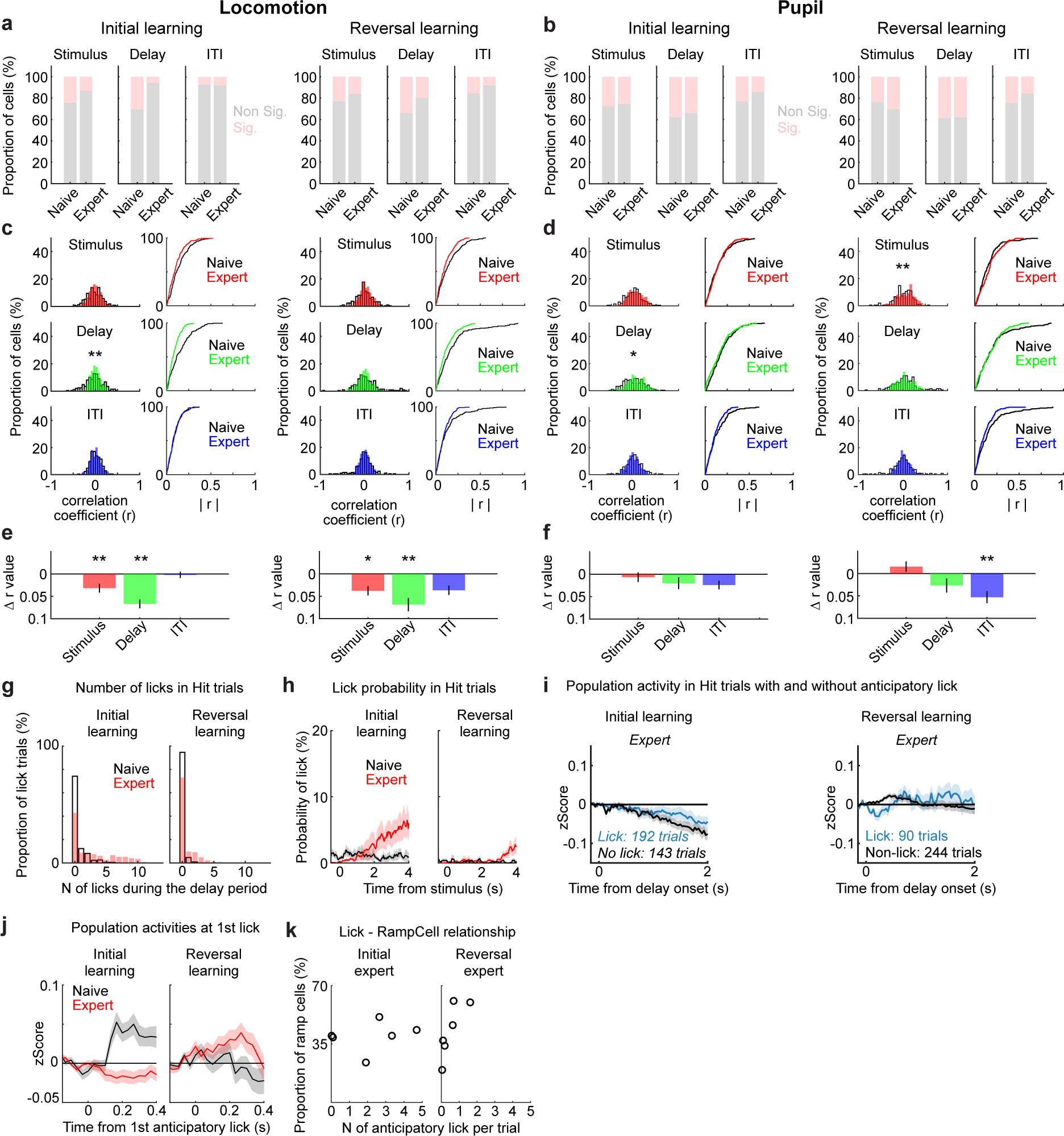
Behavioral variables are not the main source of MGB plasticity. (**a, c and e**) Correlation analysis of Ca²+ trace and locomotion in Hit trials in initial and reversal learning (N = 6 mice, n = 210 cells). The longitudinally tracked cells were used for this analysis. (**a**) Left: The proportion of the significantly correlated cells in each time window (Stimulus, Delay and ITI periods) in the initial learning (Left) and reversal learning (Right). (**c**) The distributions of the locomotion - Ca²+ correlation between the naive (black) and expert phases (color) in the initial learning (Left) and reversal learning (Right) from all cells. Top: The probability density function during the stimulus period. Middle: Probability density function during the delay period. Bottom: Probability density function during the ITI period. ITI periods is 2 s before the stimulus presentation. (**e**) The change in the magnitude of correlation. The difference of the absolute r value for each MGB neuron between the naive and expert phases (Mean ± SEM) was calculated. (**b, d and f**) Correlation analysis of Ca²+ trace and pupil size in Hit trials in the initial and reversal learning (N = 6 mice, n = 210 cells). Figure configurations are same as **a, c and e** for locomotion analysis. (**g**) The proportion of Hit trials with anticipatory licking during the delay period across all mice (N = 8 mice). The number of trials with anticipatory licking increased from the naive to expert phases in both initial and reversal learning (*p* < 0.01 for initial and reversal learning, after bonferroni correction, chi-square test). (**h**) The probability of anticipatory licking at each time bin (Mean ± SEM) in Hit trials from all mice (N = 8 mice). The probability of anticipatory lick was calculated at each time bin across Hit trials for each mouse, then averaged across all mice. (**i**) Average population activities of MGB neurons (Mean ± SEM) during the delay period in Hit trials with and without anticipatory licking across all mice (N = 8 mice). Left: Population activities in the initial expert (Lick trial = 192 trials and n = 755 cells, non-lick trial = 143 trials and n = 755 cells), Right: Population activities in the reversal expert (Lick trial = 90 trials and n = 637 cells, non-lick trial = 244 trials and n = 709 cells). (**j**) Average population activities of MGB neurons (Mean ± SEM) aligned at the first anticipatory licking onset during the delay period. Left: Average population activities of MGB neurons in the initial learning (61 trial, n = 749 cells for naive, 192 trials, n = 755 cells for expert). Right: Average population activities in the reversal learning (13 trials, n = 604 cells for naive, 90 trials, n = 637 cells for expert). (**k**) The proportion of ramping cells (ramp-up and ramp-down cells) found in k-means analysis as a function of the mean of the number of anticipatory licking during the delay period per trial in Hit trials. * and ** in the figures represent the statistical significance after Bonferroni correction. *, *p* <0.05, **, *p* <0.01.

**fig. S5.**
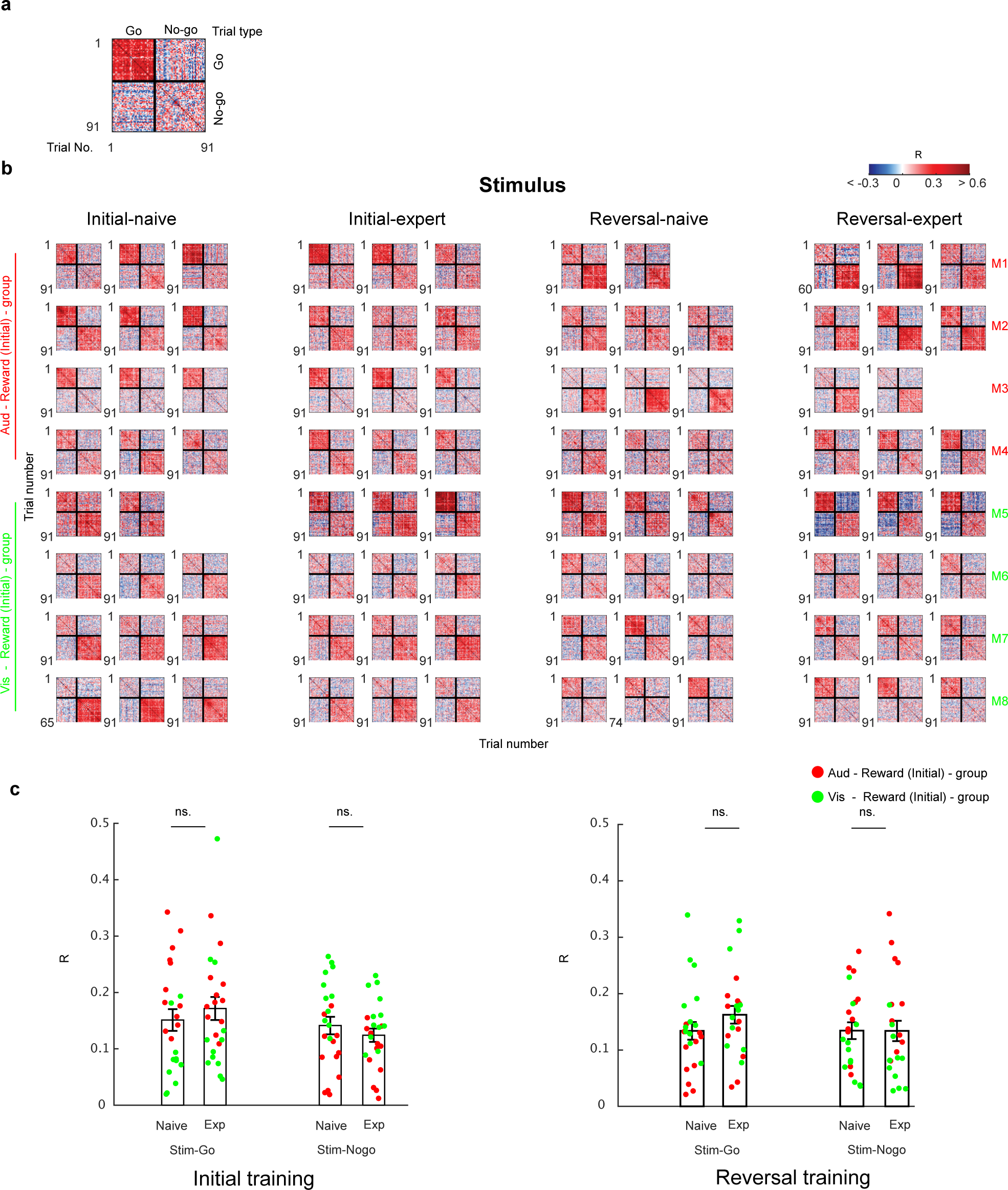
Stable evolution of MGB activity patterns during the stimulus period in the Go/Nogo learning task. (**a**) Representative trial-by-trial correlation matrix within and between Go/Nogo trials in one session. (**b**) Stimulus activity correlation matrices from all mice in different learning stages (Naive/Expert/Reversal-naive/Reversal-expert). Each matrix represents pair-wise population vector correlation (PVC) for all trials in one learning session. (**c**) Mean correlation value (R) in each session shown in (b) was grouped into initial (left) and reversal training (right). PVC in stimulus period in Expert and Reversal-expert did not significantly differ from Naive or Reversal-naive condition respectively (details in Table S1). Visualization of counterbalanced stimulus-reward pairing by color of dots. Both groups showed consistent trend of activity pattern. Red: Mice trained on auditory-reward pairing in initial training session. Green: Mice trained on visual-reward pairing in initial training session.

**fig. S6.**
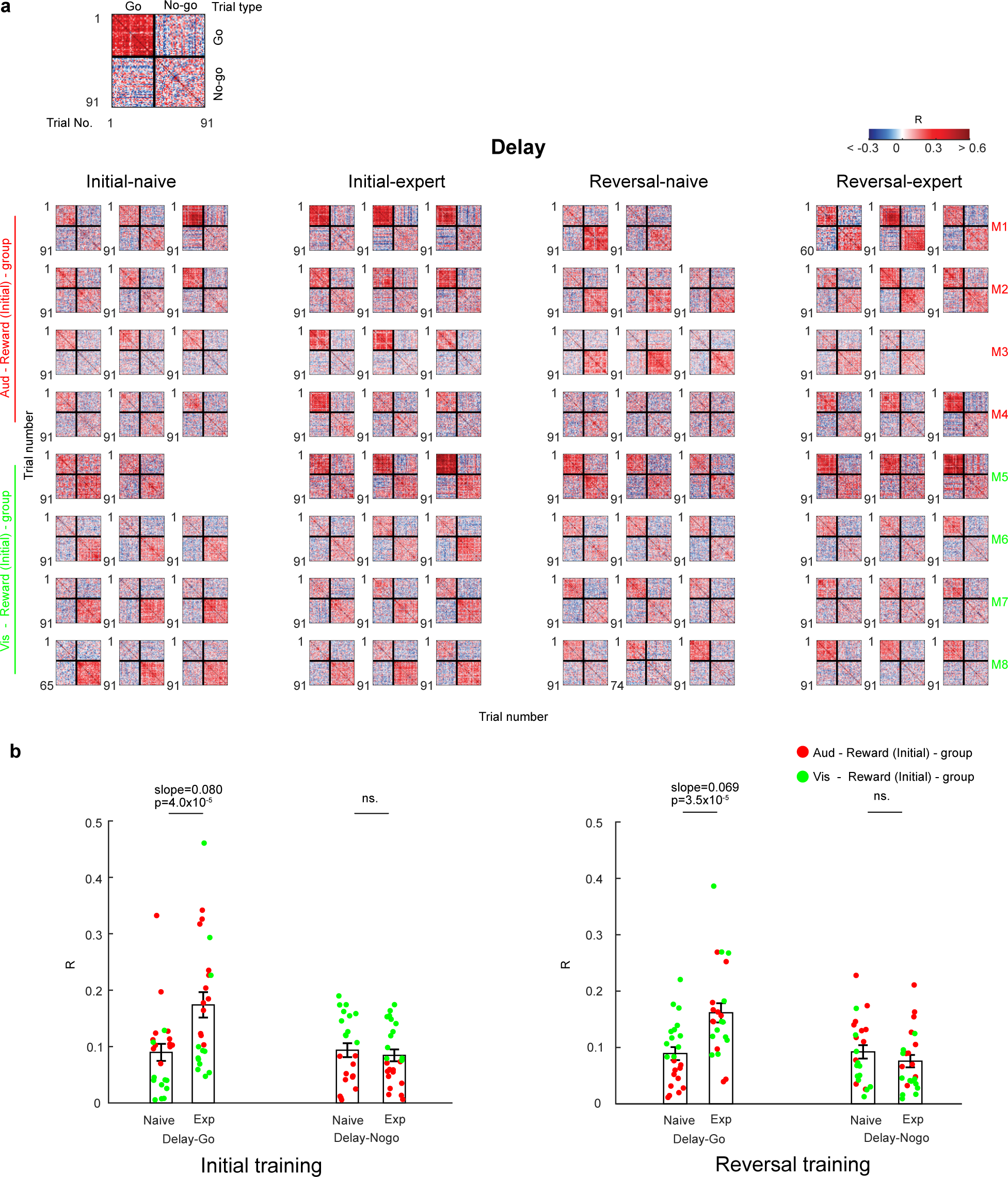
Distinct evolution of MGB activity patterns during the delay period in the Go/Nogo learning task. (**a**) Delay activity correlation matrices from all mice in different learning stages (Naive/Expert/Reversal-naive/Reversal-expert). Each matrix represents pair-wise population vector correlation (PVC) for all trials in one learning session. (**b**) Mean correlation value (R) in each session shown in (a) was grouped into initial (left) and reversal training (right). PVC in delay period in Go trial in Expert or Reversal-expert condition is higher than Naive or Reversal-naive respectively (slope > 0, *p* < 0.01, N = 8, details in Table S1). Visualization of counterbalanced stimulus-reward pairing by color of dots. Both groups showed consistent trend of activity pattern. Red dots: Mice trained on auditory-reward pairing in initial training session. Green dots: Mice trained on visual-reward pairing in initial training session.

**fig. S7.**
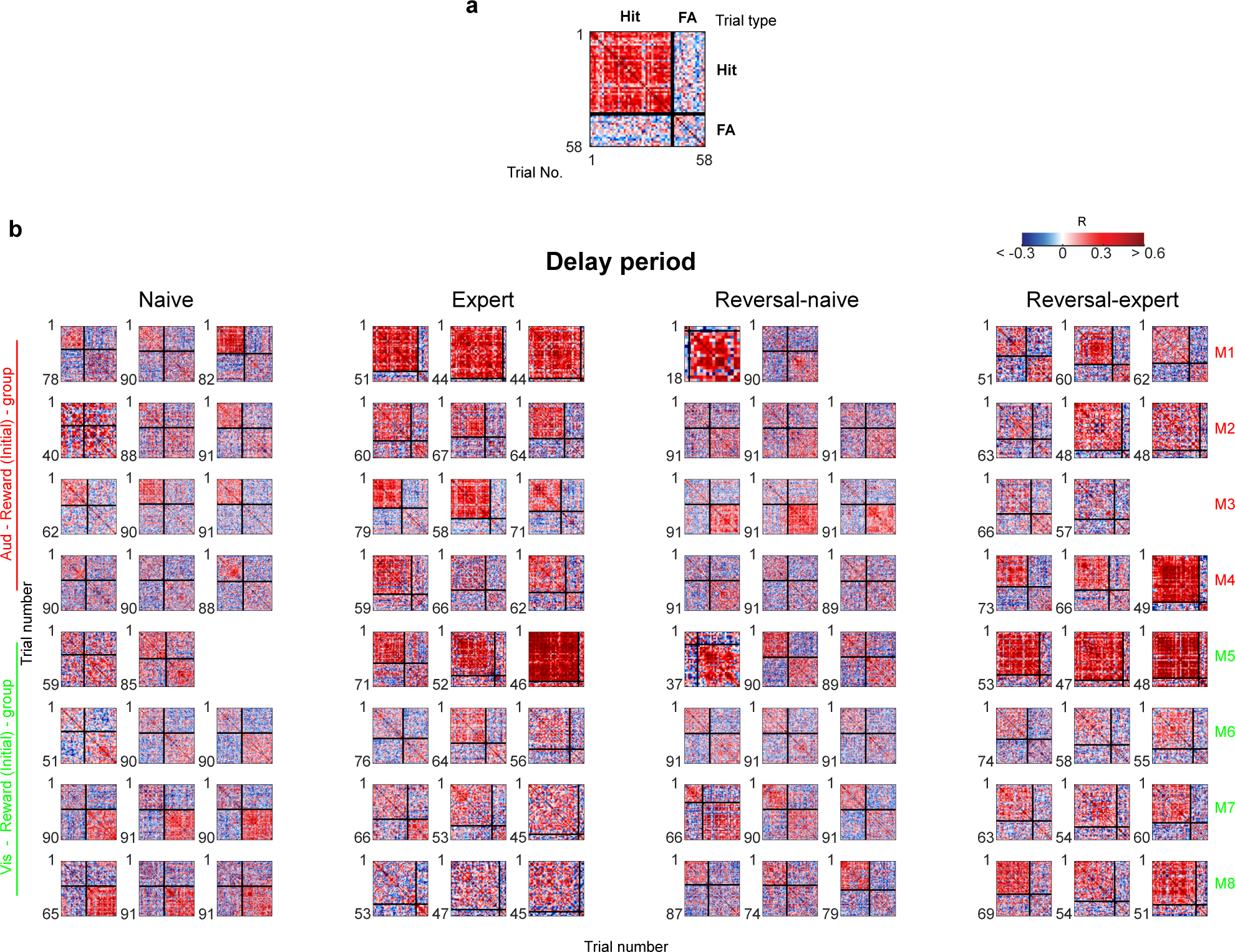
Coherence of MGB population activity is increased during the delay period of Hit but not False Alarm (FA) trials. (**a**) Representative trial-by-trial PVC matrix within and between Hit and False Alarm trials of one session. (**b**) Delay activity PVC matrices from all mice in different learning stages (Naive/Expert/Reversal-naive/Reversal-expert). Each matrix represent pair-wise PVC for all trials in one learning session. Summary data were shown in Fig.3 e, f and Table S1.

**fig. S8.**
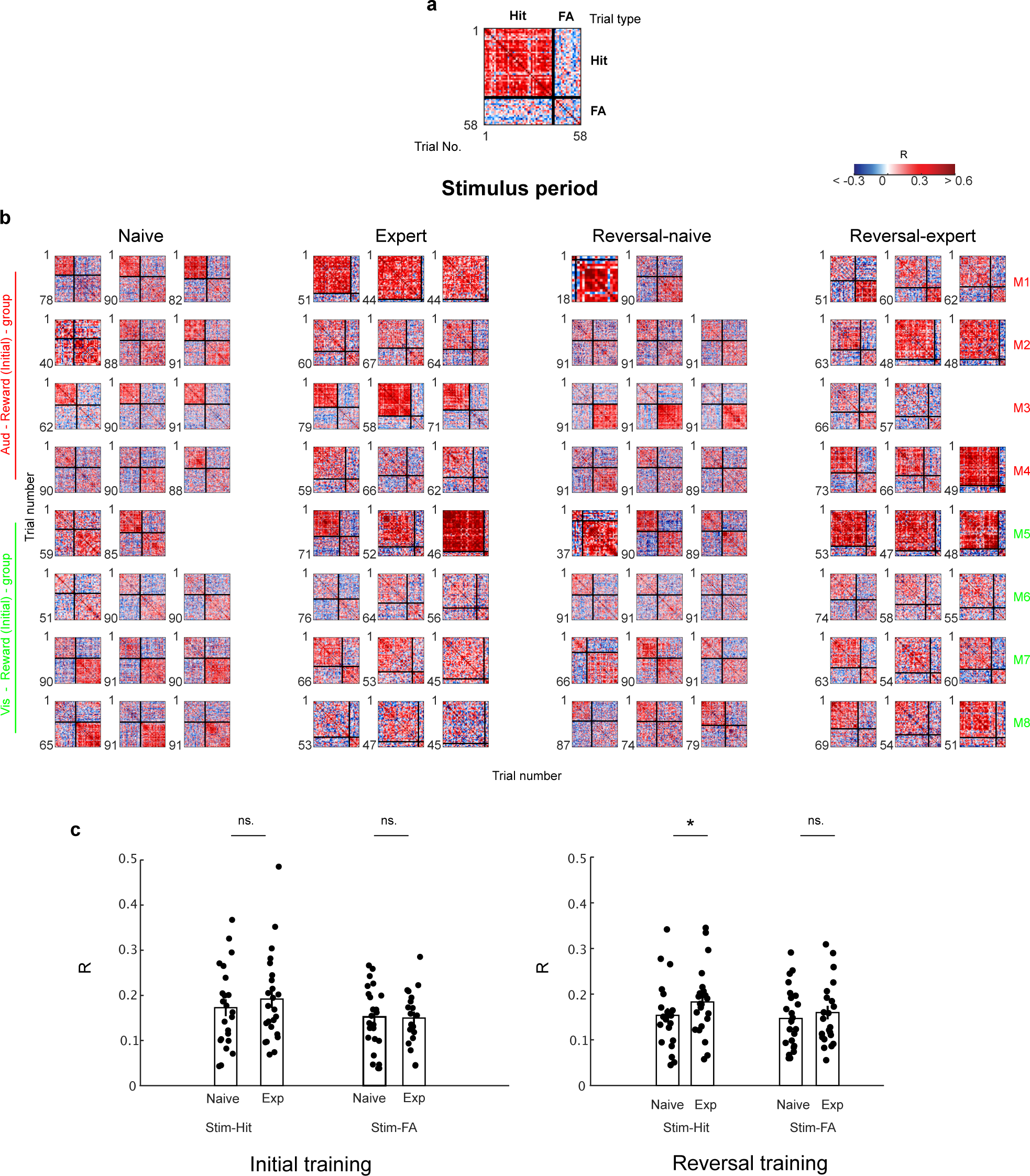
Coherence of MGB population activity is not increased during the stimulus period of Hit and False Alarm (FA) trials, but remodeled after reversal learning. (**a**) Representative trial-by-trial PVC matrix within and between Hit and False Alarm trials of one session. (**b**) Stimulus PVC matrices from all mice in different learning stages (Naive/Expert/Reversal-naive/Reversal-expert). Each matrix represent pair-wise PVC for all trials in one learning session. (**c**) Summary data. (N = 8, details in Table S1).

**fig. S9.**
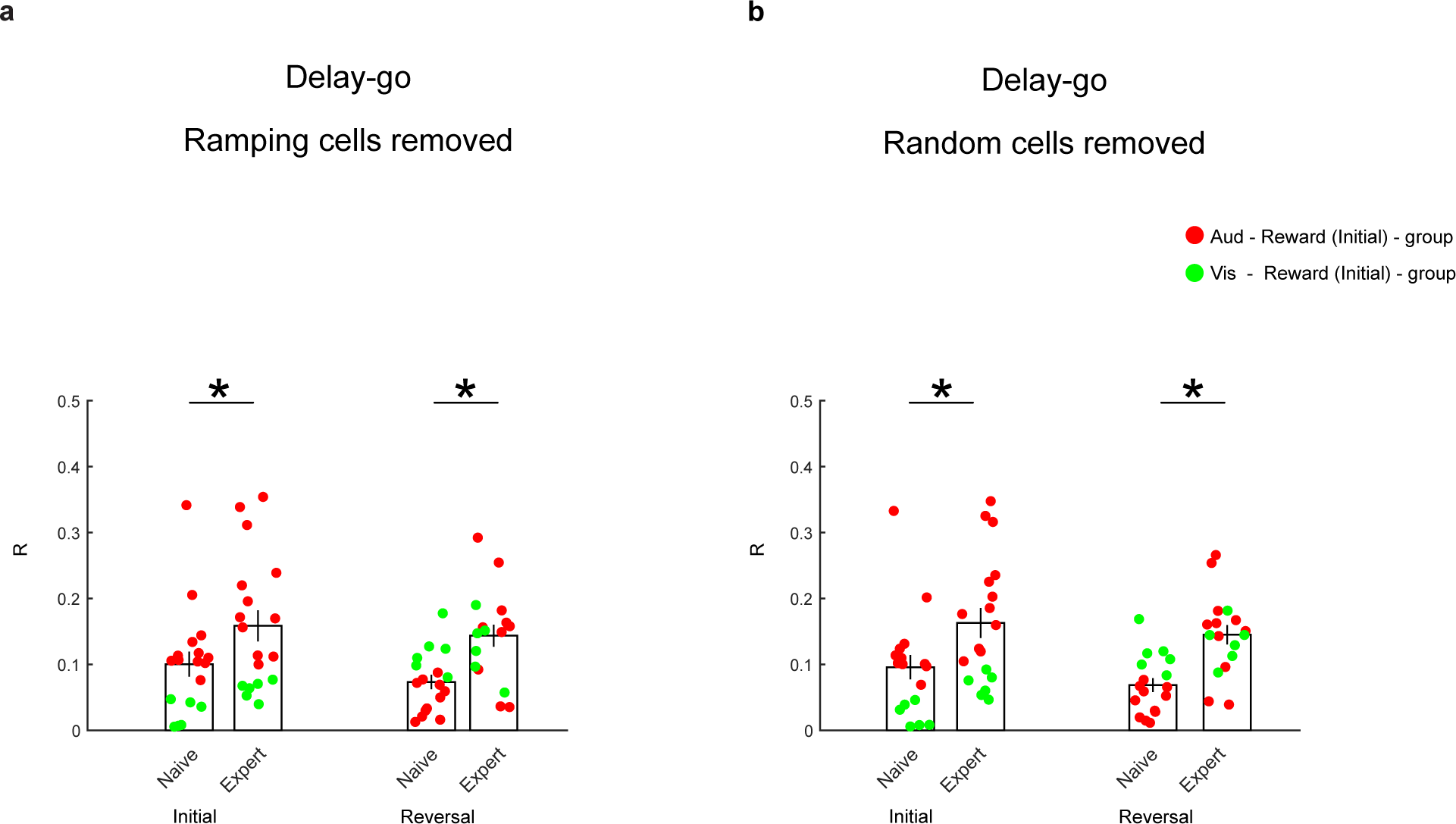
Increase of population vector correlation (PVC) in MGB during learning depends on MGB population activity and is not driven by ramping-cells. (**a**) Mean trial-by-trial PVC (R) during the delay period of Go trials in different learning stages. Increase of PVC persisted even if ramping cells (ramp-up and ramp-down, Fig. 2a, b) were removed (* *p* < 0.01, details in Table S1). (**b**) Increase of PVC was dependent on the MGB population activity instead of subsets of neurons. Same number of cell as ramping cells (ramp-up + ramp-down) in each mouse were randomly removed before calculating PVC (number of random cells =30, \**p* < 0.01, details in Table S1).

**fig. S10.**
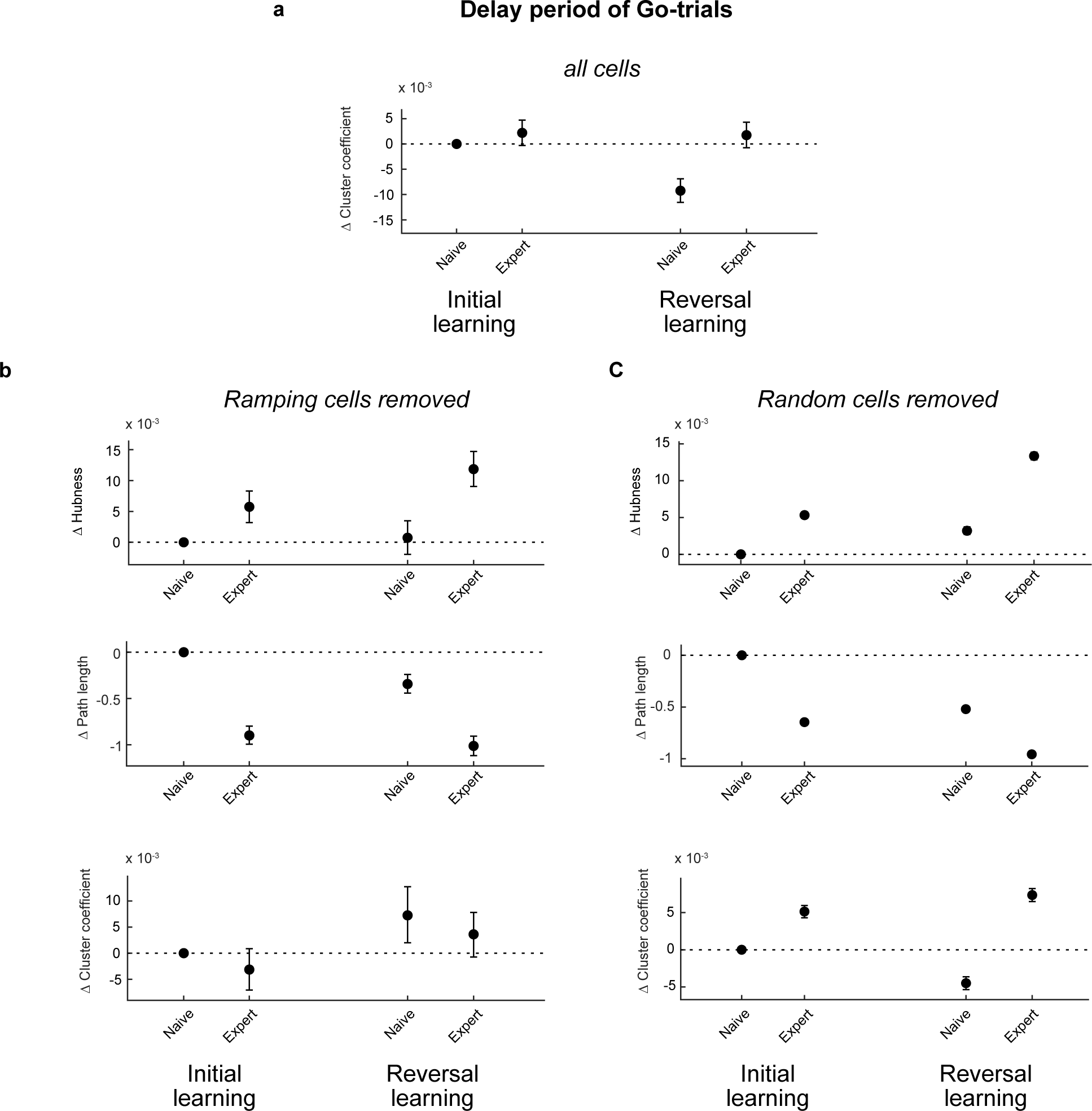
MGB co-activity network structure does not depend on ramping cells. (**a**) Decrease of local connectivity was observed specifically in reversal-naive learning stage when reward contingency was changed, suggesting a re-organization of pre-configured connectivity that was established in Initial-expert stage. (**b**) Ramp-cells were not the sole contributors of connectivity re-organization. Removing ramp-cells (ramp-up and ramp-down, Fig. 2a, b) or random cells did not abolish the change of network architecture which was an indication of MGB general ensemble remodeling during reward-association learning task (detail in Methods).

**fig. S11.**
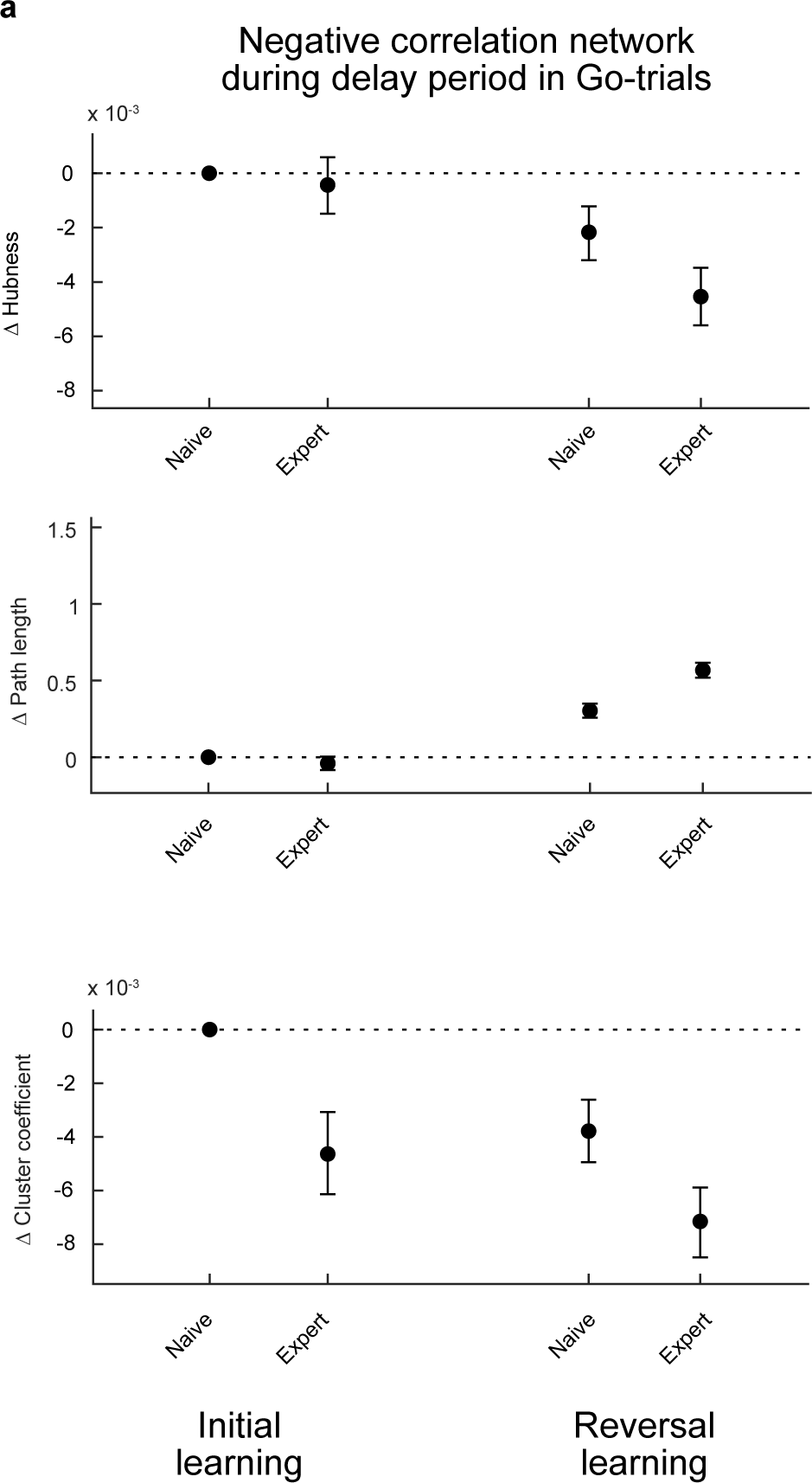
Negative correlation co-activity network during delay period in Go-trials. (**a**) There was no prominent change of global negative co-activity structure from Naive to Expert stage in Initial and Reversal learning phase which was in accordance to the increase of global positive co-activity network (detail in Methods). Local negative co-activity network became less connected from Naive to Expert stage suggesting an increase of connectivity during delay period when the mouse learned the task structure. Error bars: 95 % confidence interval of mean.

**fig. S12.**
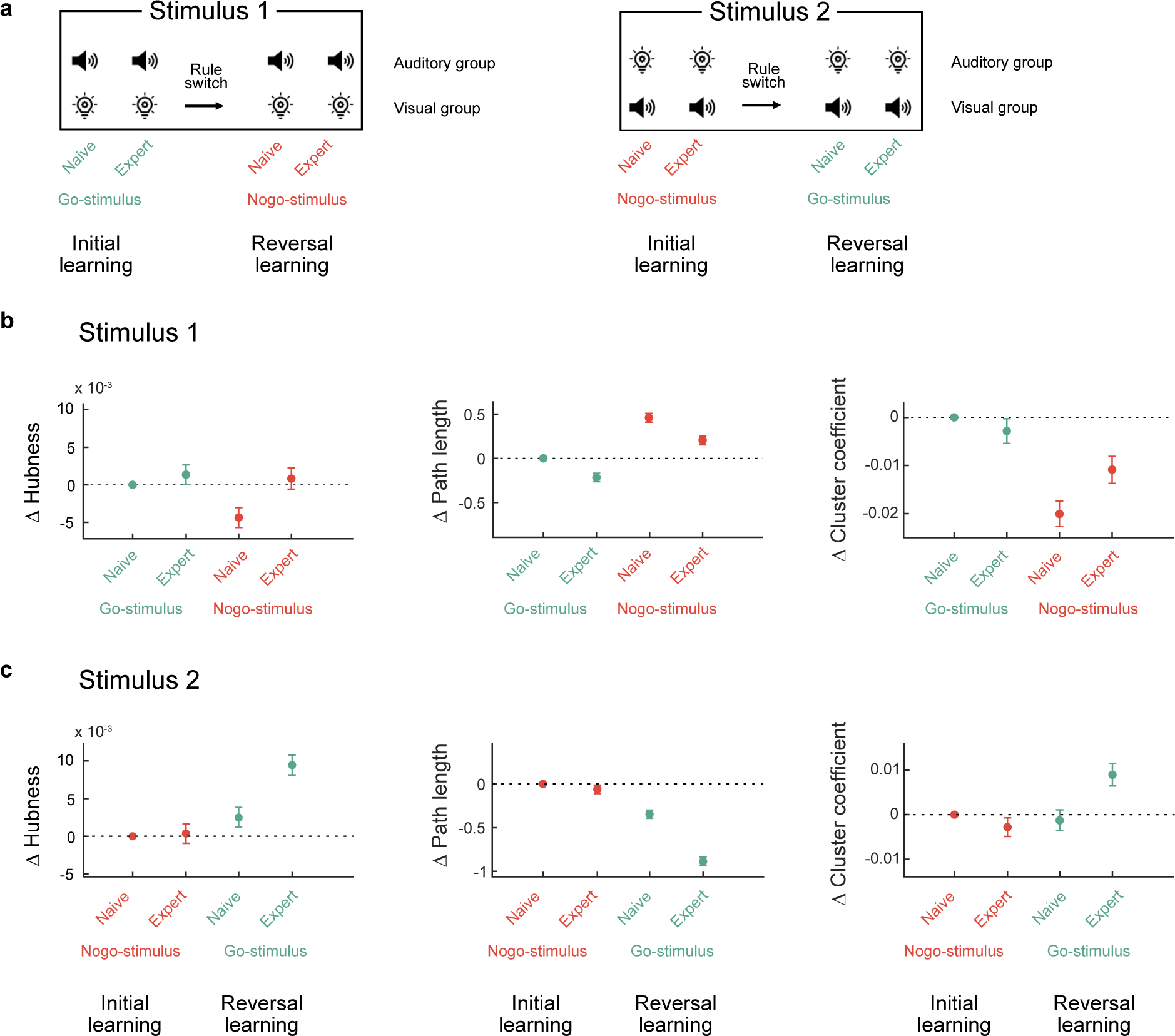
MGB co-activity networks during the stimulus period. (**a**) Experimental design. (**b** and **c**) Change of hubness, path length and cluster coefficient across all neuron pairs in relation to the naive Go (or) Nogo condition (detail in Materials and Methods). There was no obvious change of global and local connectivity in initial training phase during stimulus presentation period for within group identical stimulus 1 and 2. Conversely in reversal learning, global and local connectivity of both Go and Nogo stimuli were strengthened. Neural representation of stimuli that were associated or dissociated with reward contingency were remodeled in the reversal naive phase. Error bars: 95 % confidence interval of mean.

**fig. S13.**
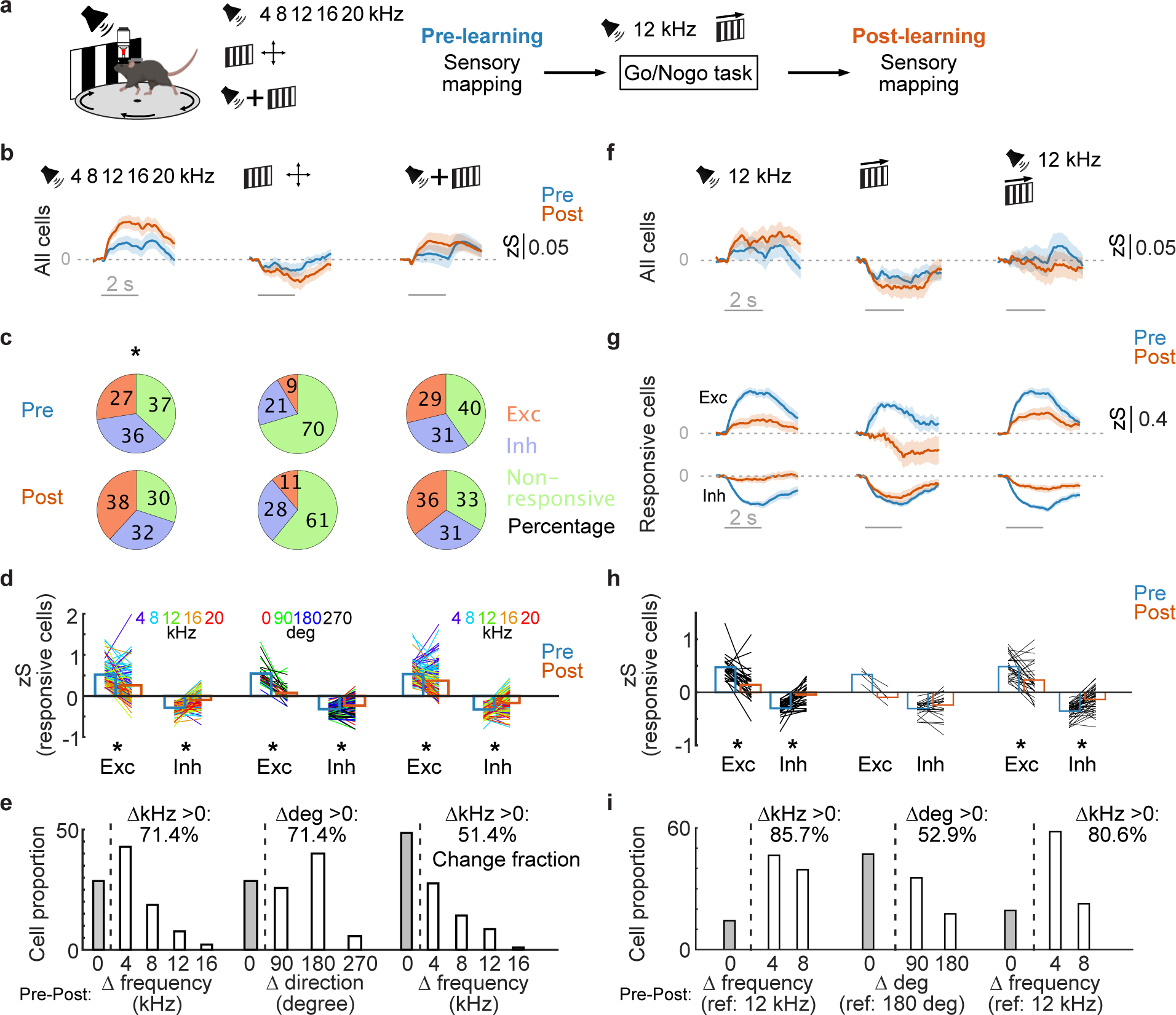
Sensory coding in MGB is not biased to the reward-associated stimuli in non-rewarded sensory mapping sessions. (**a**) Schematics of sensory mapping. Various auditory and visual stimuli were presented as unisensory stimulus, and the combination of them were presented as multisensory stimulus. (**b**) Average population calcium activity in uni- and multi-sensory trials before and after learning (n = 233 cells, 6 mice). (**c**) Proportions of the excited and inhibited cells to auditory, visual and multi-sensory stimuli. (**d**) Transition of the sensory response amplitude from the cells showing the significant response at pre-learning stage were plotted. Each line represents sensory responses before and after learning from the same cell for a specific stimulus feature (tone frequency or grating direction). (**e**) Changes of the best frequency/direction tuning across learning. (**f**) Averaged population calcium activity to the reward-conditioned sensory stimuli (12 kHz tone, rightward grating, combination of them, n = 233 cells, 6 mice). (**g**) Average population responses of the excited or inhibited cells to the conditioned stimuli. (**h**) Transition of the sensory response amplitude to the conditioned stimuli. Each line represents the sensory response of the same cell before and after learning. (**I**) Changes of the best frequency/direction tuning referred to the conditioned stimuli. Statistical details in Table S1. Parts of panel S13a were created with BioRender.com.

**fig. S14.**
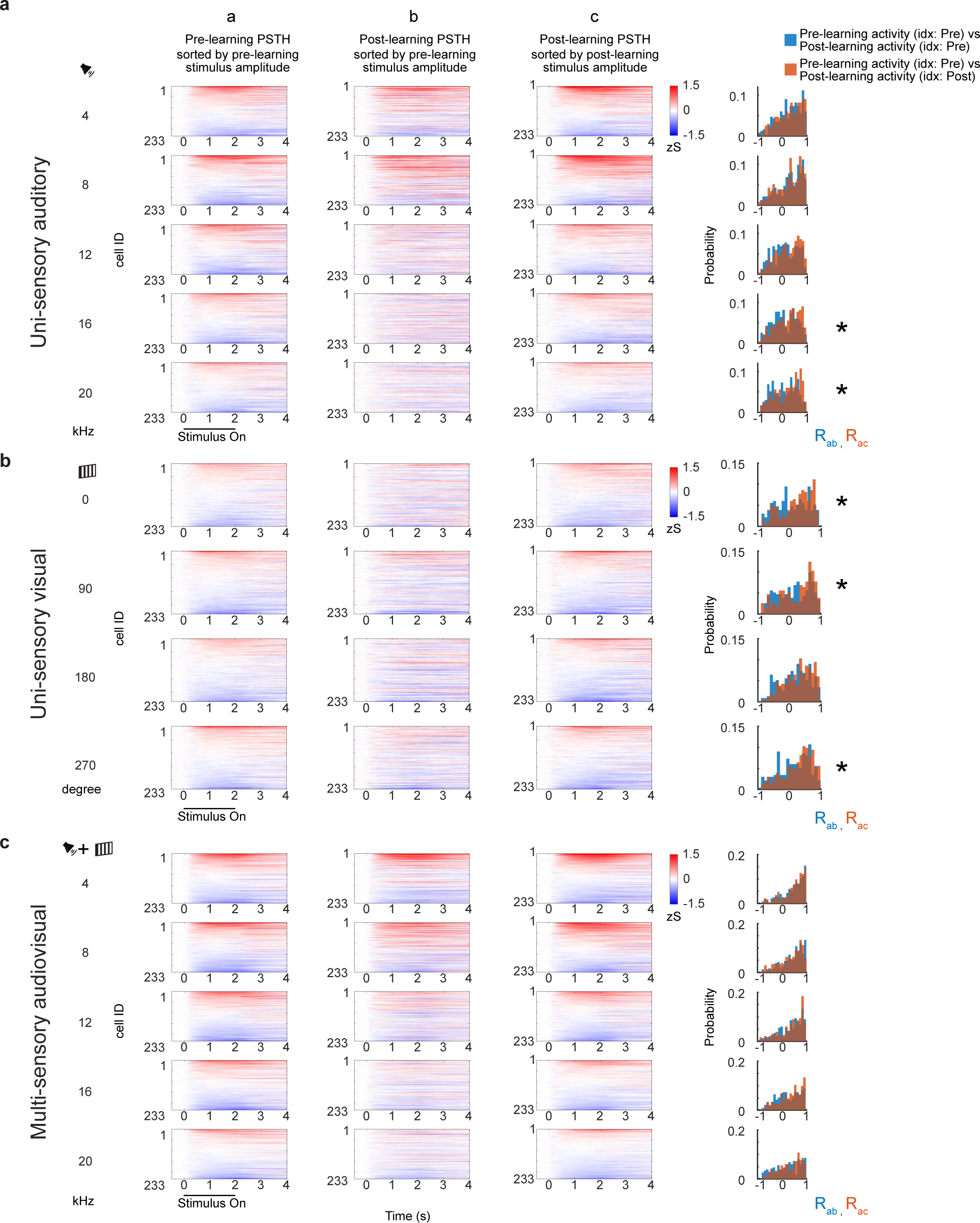
MGB single cell responses across sensory mapping. (**a**) Heat-map shows average peri-stimulus time histogram (PSTH) of cell response in pre-learning for uni-sensory auditory stimulus sorted by pre-learning response amplitude during stimulus presentation (column a). Same sorting index was used to sort post-learning PSTH (column b). Reduced response amplitude with same sorting index indicated a remapping of sensory response to the same auditory frequency. Post-learning responses were not diminished after learning if cells were sorted with amplitude in post-learning session (column c). Correlation of cell response time series using two different indexes was compared to quantify the extent of remapping (R_ab_:R_ac_, 2-sample ks-test). (**b, c**) Same measurement for uni-sensory visual response and multi-sensory response.

**fig. S15.**
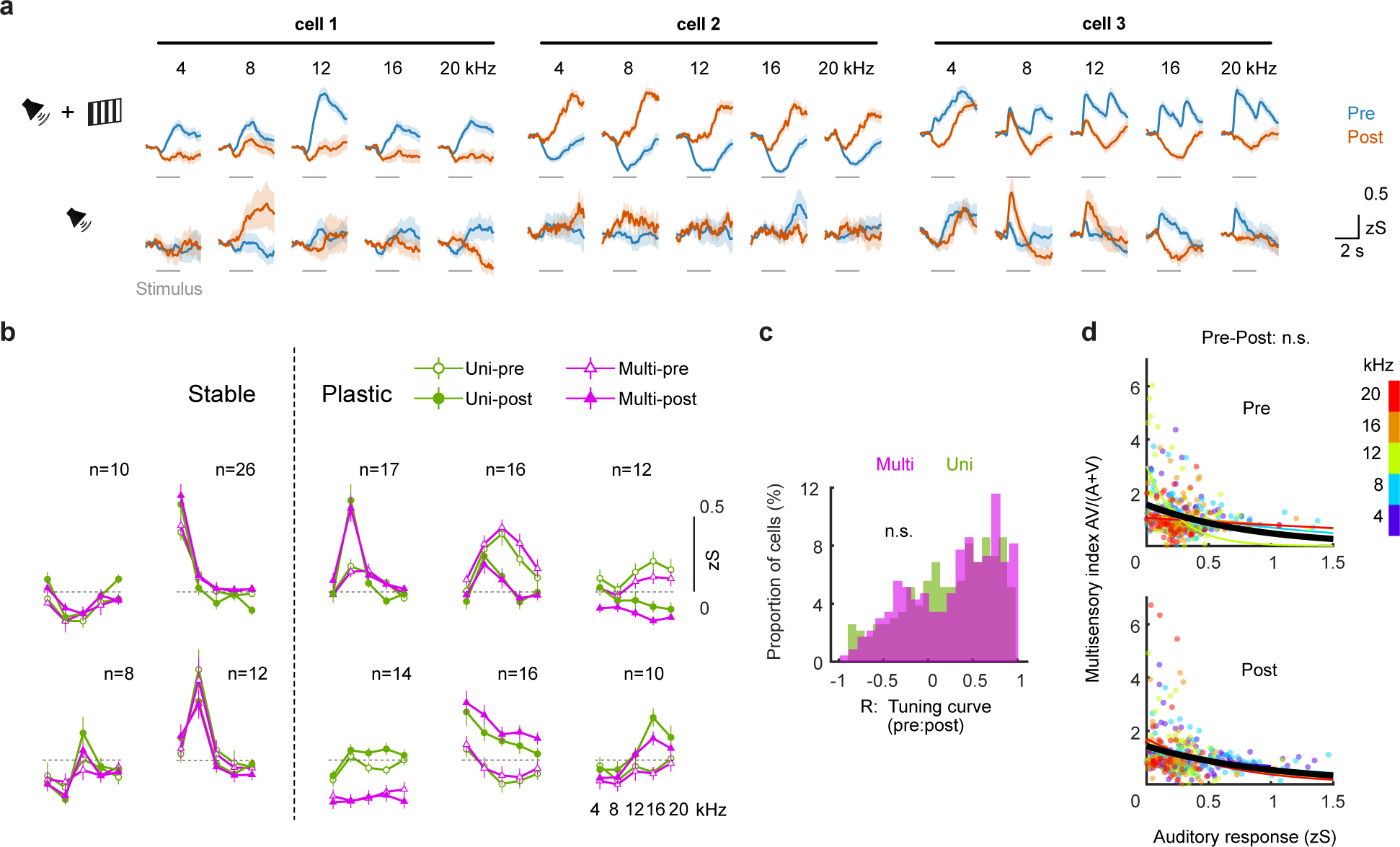
Single cell tuning properties are not biased to the reward-associated stimuli in non-rewarded sensory mapping sessions. (**a**) Example sensory responses of individual MGB neurons to auditory and multisensory stimuli. Individual neurons showed stable or plastic sensory response patterns to auditory and multisensory stimuli. (**b**) Transition of tone-frequency tuning across learning. The neurons showing a similar changing pattern of frequency tuning were grouped by k-means clustering. (**c**) Distribution of tuning curve similarity of individual neurons across learning. Tuning curve similarities are comparable in auditory and multisensory trials. R is Pearson’s correlation coefficient between pre- and post-learning tuning curve (n = 233 cells, 6 mice). (**d**) Multisensory index as a function of unisensory tone response. Two-dimension distribution of unisensory response (auditory) versus multisensory index was compared between pre-learning and post-learning conditions (5 frequencies were pooled, *p* > 0.05, details in Table S1).

**fig. S16.**
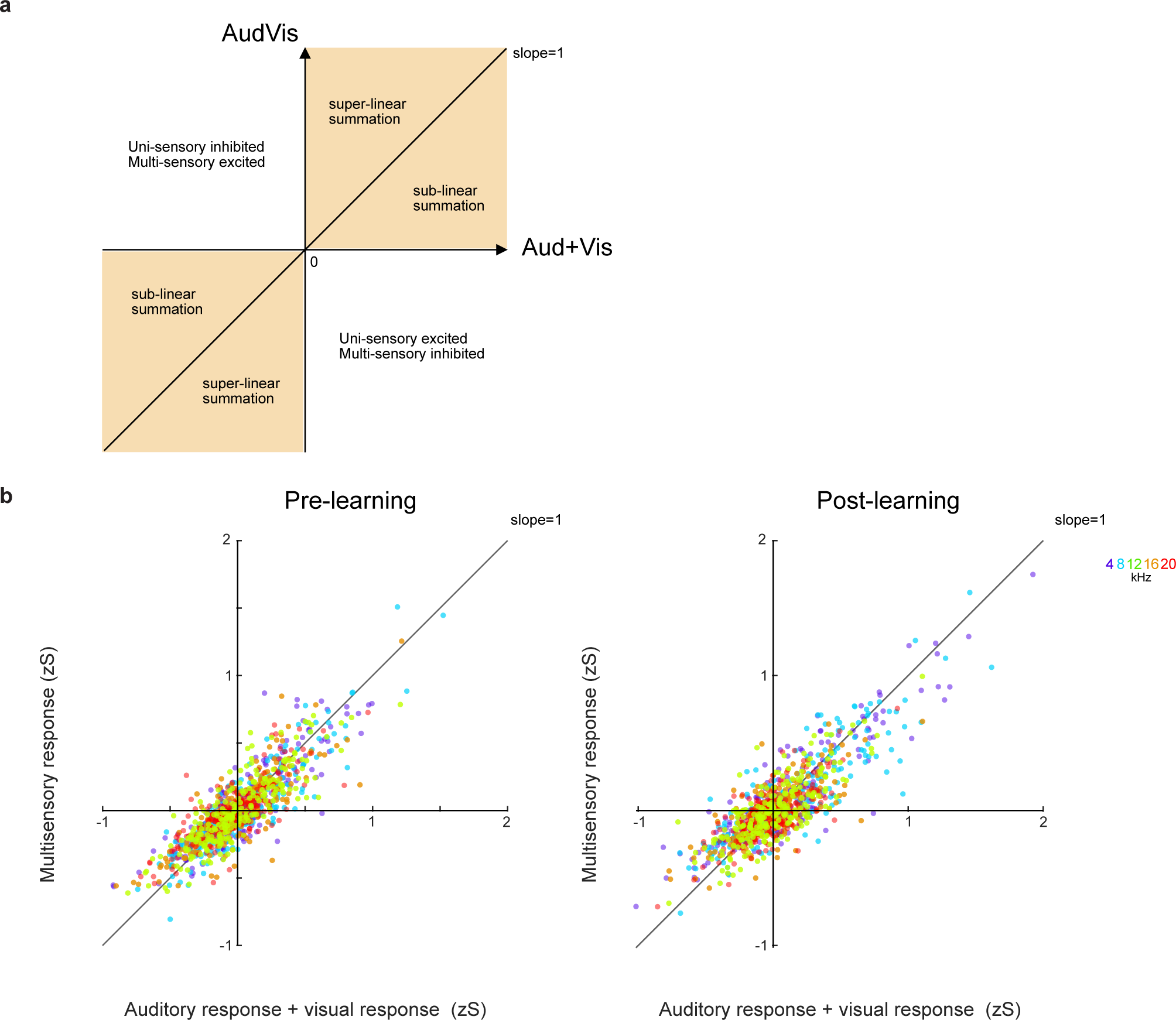
Multisensory integration remains unchanged after cross-modal reward learning. (**a**) Multi-sensory integration principle is described in each quadrant. Data in 1st and 3rd quadrant were included in analysis in fig. S15d. (**b**) Average response of Multi-sensory trials (Audiovisual trials) for each cell and for each auditory frequency were plotted against the sum of average response of Uni-sensory trials (Auditory and Visual trials). Diagonal line represents unity between multisensory responses and the linear sum of individual unisensory responses. Response distribution was not different between pre-learning and post-learning sessions (2-dimensional 2-sample KS-test, details in Table S1). Each dot indicates a cell (repeat measurement form 1-3 sessions).

**fig. S17.**
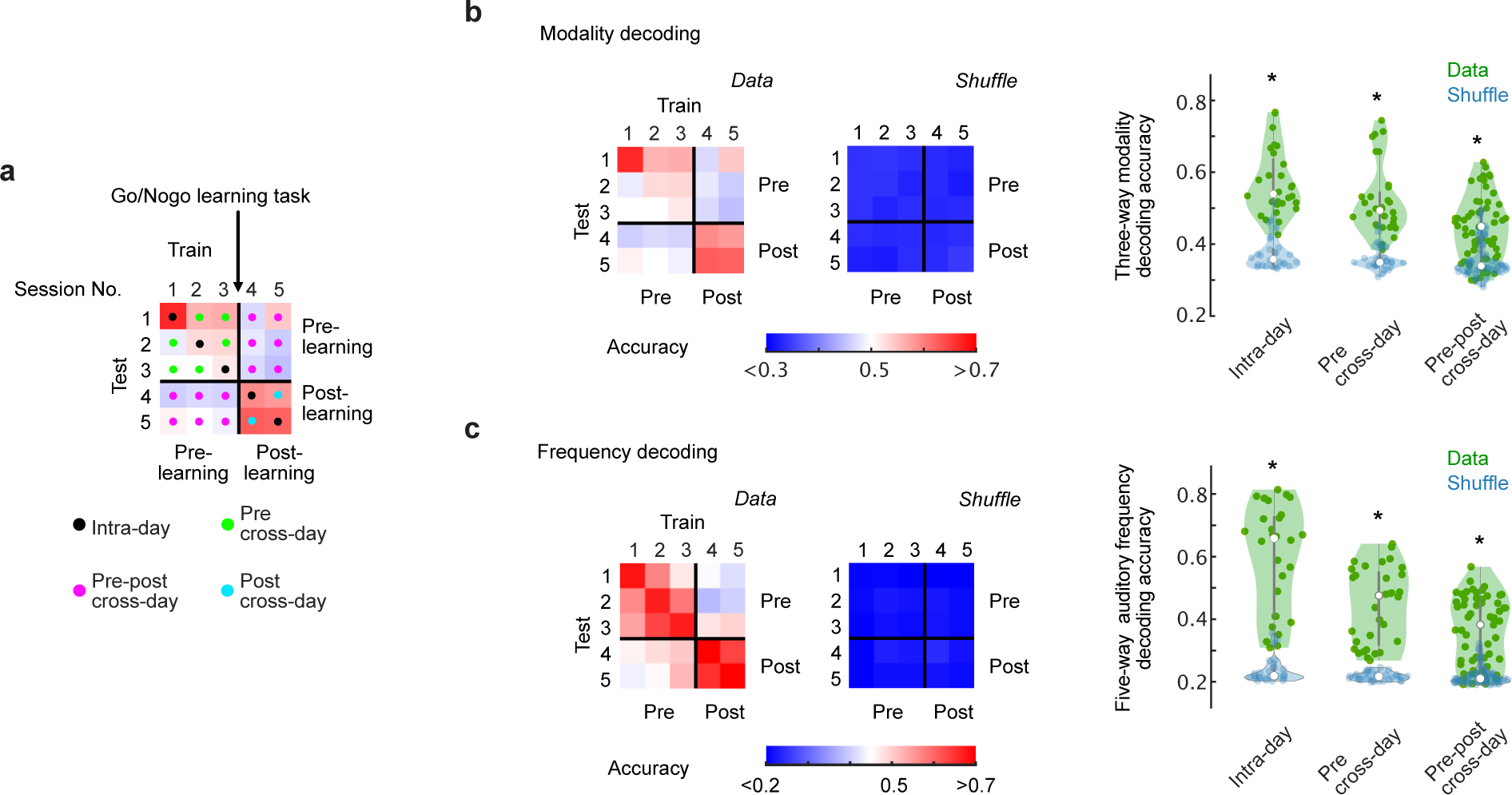
The total MGB population reliably encodes sensory information. (**a**) Decoding design matrix. Linear-SVM decoder was trained and tested in a session-by-session manner. (**b**) Three-way modality decoding. Left: Matrix of modality decoding accuracy from one representative mouse. Middle: Matrix of modality decoding accuracy from the same mouse generated by the shuffled data. Right: Decoding accuracies grouped by intra-day, pre cross-day and pre-post cross-day conditions. Modality decoding accuracies were above chance (\**p* < 0.01). Each dot represents the one session data from one mouse. Shuffle data showed the chance level decoding accuracy. (**c**) Five-way tone-frequency decoding. The figure structures are the same as (b). Frequency decoding accuracies were above chance (\**p* < 0.01, details in Table S1).

## Supplementary Tables

**Table. S1.**
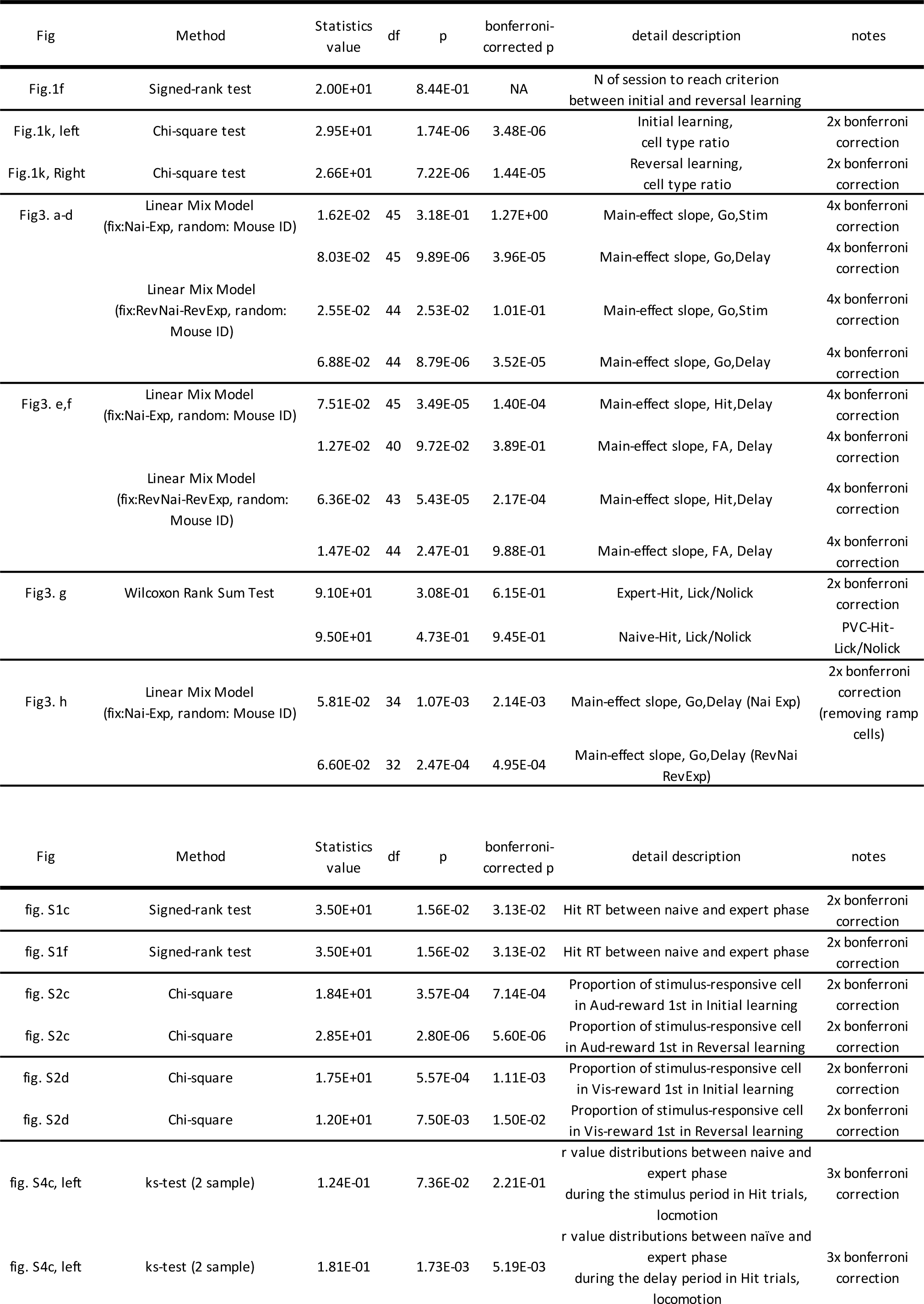
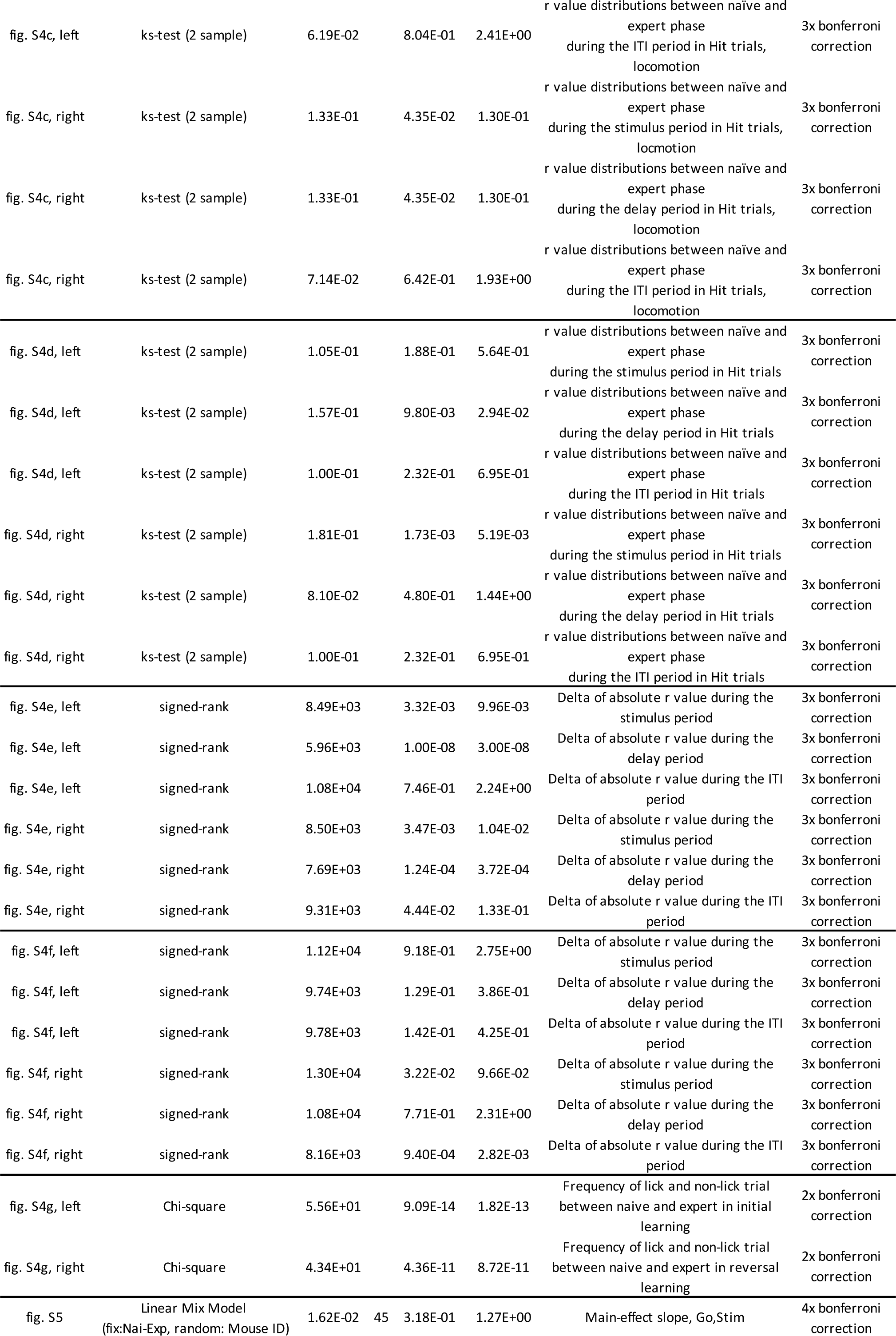
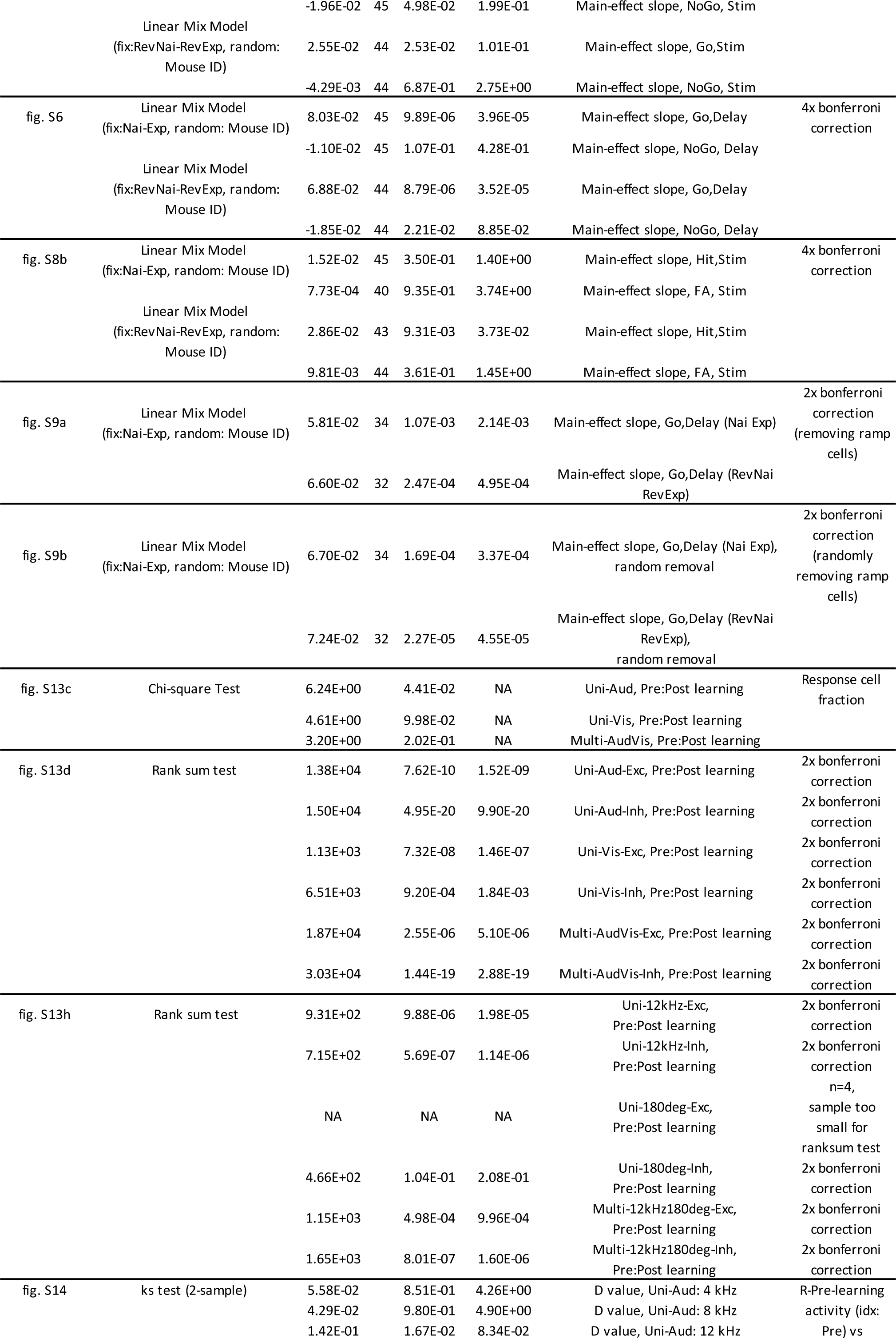
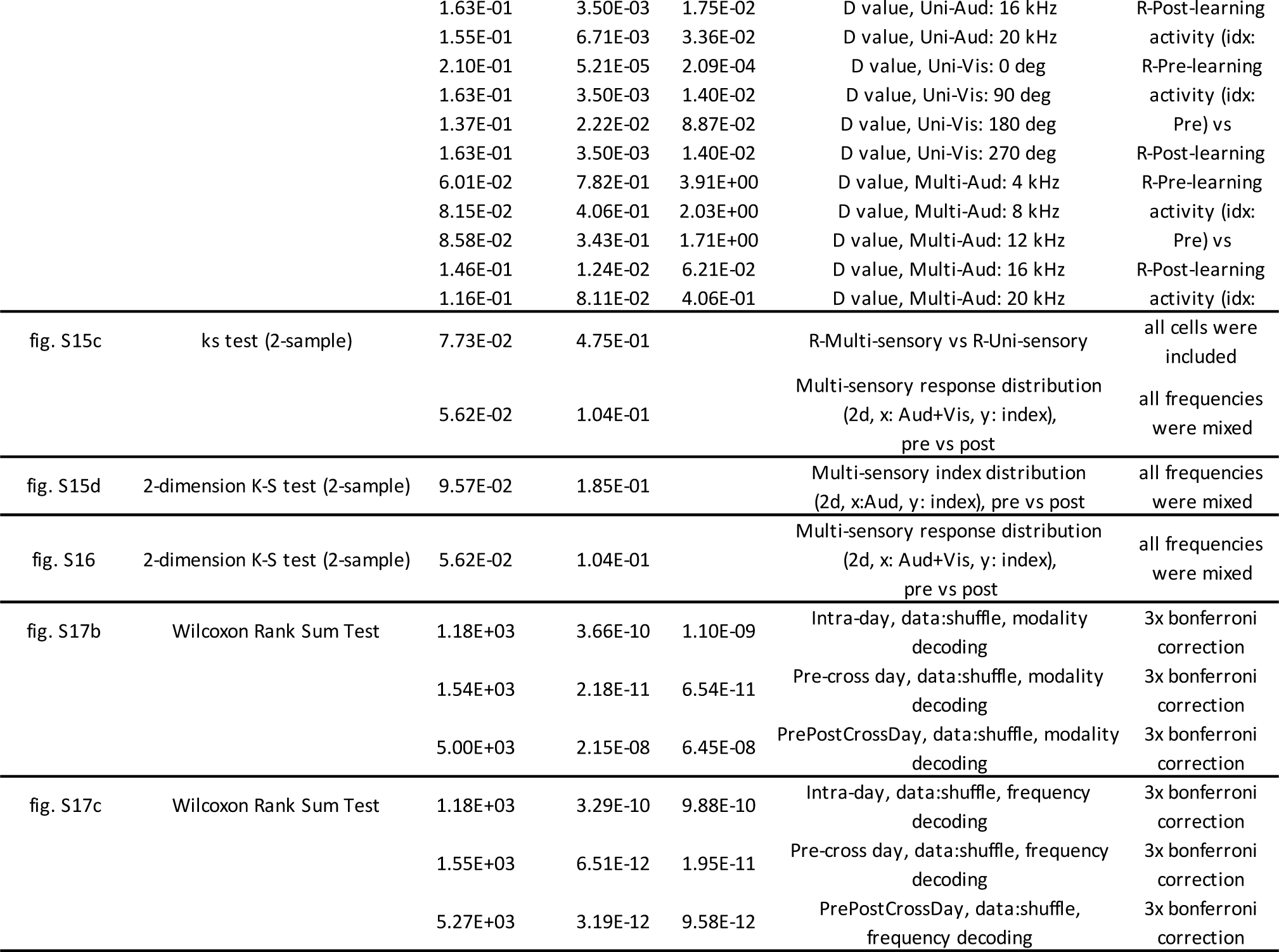
Summary of statistical tests.

